# COVID-19 ORF3a Viroporin Influenced Common and Unique Cellular Signalling Cascades in Lung, Heart and Brain Choroid Plexus Organoids with Additional Enriched MicroRNA Network Analyses for Lung and Brain Tissues

**DOI:** 10.1101/2023.09.20.558551

**Authors:** Soura Chakraborty, Shrabonti Chatterjee, Subhashree Mardi, Joydeep Mahata, Suneel Kateriya, Pradeep Punnakkal, Gireesh Anirudhan

## Abstract

Tissue specific implications of SARS-CoV-2 encoded accessory proteins are not fully understood. SARS-CoV-2 infection can severely affect three major organs – the heart, lung, and brain. We analysed SARS-CoV-2 ORF3a interacting host proteins in these three major organs. Further we identified common and unique interacting host proteins, their targeting miRNAs (lung and brain), and delineated associated biological processes reanalysing RNA-seq data from the brain (COVID-19 infected/uninfected Choroid Plexus Organoids study), lung tissue from COVID-19 patients/healthy subjects, and cardiomyocyte cells based transcriptomics analyses. Our *in silico* studies showed ORF3a interacting proteins could vary depending upon tissues. Number of unique ORF3a interacting proteins in brain, lung and heart were 10, 7 and 1 respectively. Though common pathways influenced by SARS-CoV-2 infection were more, unique 21 brain and 7 heart pathways were found. One unique pathway for heart was negative regulation of calcium ion transport. Reported observations of COVID-19 patients with the history of hypertension taking calcium channel blockers (CCBs) or dihydorpyridine CCBs had elevated rate of intubation or increased rate of intubation/death respectively. Also likelihood of hospitalization of chronic CCB users with COVID-19 was more in comparison to long term Angiotensin Converting Enzyme inhibitors/Angiotensin Receptor Blockers users. Further studies are necessary to confirm this. miRNA analysis of ORF3a interacting proteins in brain and lung revealed, 2 of 37 brain miRNAs and 1 of 25 lung miRNAs with high degree and betweenness indicating their significance as hubs in the interaction network. Our study could help in identifying potential tissue specific COVID-19 drug/drug repurposing targets.

## 1. **Introduction**

Risk of SARS-CoV-2 (COVID 19) that caused the recent pandemic is far from being over because of the lack of complete protection by the current vaccines and ever emerging variants. This situation is more alarming by the detection of COVID19 antibodies in a wide variety of animals both domestic and wild as well as the detection of COVID19 infection itself, generating a possibility of these animals becoming potential as well as observed reservoir hosts. ^1–5^ In addition, observation of unique COVID19 clades specific to mink as well as variations observed among different strains^6^ suggests the emergence of new variants independently in these hosts that could pose an increased risk to humans. Also, the lack of effective treatment targeting COVID19 is a major lacuna that impairs our combat against it including development of long COVID. COVID 19, with a positive strand RNA as its genetic material is a member of Coronavirus family that cause many diseases in vertebrates including humans. In 2002, *S*evere *A*cute *R*espiratory *S*yndrome (SARS-CoV) identified in humans with mortality rate of 15 % all over the world^7^ and *M*iddle *E*ast *R*espiratory *S*yndrome (MERS) in 2012 with a mortality rate of 35%^8^ are coronaviruses.

Initially, SARS-CoV-2 pathogenesis was focused on respiratory pathologies, mainly on the symptoms including cough, fever, common cold and respiratory distress. However, recent evidences have shown that SARS-CoV-2 is not only confined to the organs of the respiratory system, but also invades other organs such as the brain, heart, kidney, intestine etc.^9–13^ For the development of an effective treatment against COVID19, investigating the mechanism of action of different viral encoded protein is paramount. This is envisaged in the use of Spike protein as a vaccine candidate, as well as testing putative drug molecules targeting RNA-dependent RNA polymerase (RdRp), viral proteases 3CLpro and PLpro, etc.^14–19^

Viroporins are small channel forming integral membrane viral proteins present in many different viruses (reviewed in Scott and Griffin).^20^ They vary in their number of amino acids, transmembrane domains (1 – 3), ion selectivity (H ^+^, Na^+^, K^+^, Ca^2+^ and Cl^-^) and have diverse functions depending upon the virus family they belong (reviewed in Scott and Griffin).^20^ Diverse functional roles of viroporins in different viral families are observed include aiding in cell entry, cell lysis, particle production, viral spread, as TNF antagonist, influencing and manifesting pathogenesis, and mitochondrial permeability (reviewed in Scott and Griffin).^20^ Adamantane inhibitors, used in the treatment of Influenza A virus, targeting M2 protein^21–25^ is halted because of resistant polymorphisms present in majority of circulating strains (reviewed in Scott and Griffin).^20^ Another viroporin targeted was p7 protein of of HCV (Hepatitis C virus), where the inhibitors fall in three categories viz., adamantanes, alkyl imino-sugars and HMA (hexamethylene amiloride). Recombinant protein or peptides were used in vitro for their identification.^20,26–28^ Though genotype-dependent resistance was observed,^29,30^ broad-spectrum ligands targeting multiple targets could be explored as an option to overcome this resistance.^31^ Warranting more studies, inhibition of Dengue virus replication by amantadine was reported by an in vitro study^32^ and administration of amantadine at onset and 2 to 6 days after onset was observed to be diminishing the symptoms of Dengue infection in comparison to control group.^33^ Hence looking for viroporin interacting proteins and different pathways would be an alternate strategy for therapeutic intervention.

Viroporins associated with Coronaviruses are E, 3a, ORF8a and ORF4a.^20,34^ In SARS-CoV-2, ORF3a is one of the putative viroporins. When purified from heterologous expression systems it is reported to exist as a 62-kDa dimer and 124-kDa tetramer with each protomer having 3 transmembrane domains.^35^ ORF3a induce inflammatory response in the host^36–39^ and also mediates optimal replication.^40^ Retrospective analyses of 16 studies for the levels of inflammatory markers like C-reactive protein (CRP), procalcitonin (PCT), serum ferritin, erythrocyte sedimentation rate (ESR) and interleukin-6 (IL-6) showed positive correlation with COVID-19 severity.^41^ Antibody response is observed against SARS-CoV ORF3a^42,43^ and SARS-CoV-2 ORF3a.^44,45^ Recently, the role of viroporin ORF3a in the induction of NLRP3 mediated inflammatory responses has also been reported.^36,39^ It is also observed that SARS-CoV-2 ORF3a viroporin has relatively weaker pro-apoptotic activity compared to ORF3a viroporin of SARS-CoV when expressed in cell lines.^46^ It is also speculated that relatively mild infection of the SARS-CoV-2 might give it an advantage in spreading.^46^ Observation of SARS-CoV-2 ORF3a unlike SARS-CoV ORF3a promoting lysosomal exocytosis-mediated viral egress,^47^ blocking autolysosome formation by interfering with the assembly of STX17-SNAP29-VAMP8 SNARE complex,^48,49^ and mutation in ORF3a associated with increased mortality rate in SARS-CoV-2 infection increases its importance.^50^

This warrants the investigation of ORF3a and(or) other interacting partners involved in various pathways as plausible drug target(s). The targeted disruption at interface of interaction of viroporin and host protein might also be a useful strategy to overcome drug resistant phenomenon. In our *in silico* analyses reported here, we attempted to find out various pathways where SARS-CoV-2 ORF3a and its interacting partners are involved. Common and unique pathways in lung, heart and brain choroid plexus organoids were found in addition to the observation that SARS-CoV-2 ORF3a interacting partners getting regulated after SARS-CoV-2 infection. We were able to find 10, 7 and 1 unique interacting proteins out of 200, 197 and 175 interacting proteins that were regulated in brain choroid plexus organoids, lung and heart respectively after SARS-CoV-2 infection. Looking for probable biological processes of brain and lung regulated by miRNAs, we analysed miRNet for interacting miRNAs of 8 proteins in these tissues that interact with ORF3a. We could find 2 out of 37 miRNAs in brain and 1 out of 25 miRNAs in lung with high degree and betweeness that signifies the role of these miRNAs as hubs. We also looked for SARS-CoV-2 influenced miRNAs as well as proteins that interact with ORF3a interacting proteins in a tissue specific manner in brain and lung to find out prominent biological processes in these tissues.

## 2. **Materials and methods**

### 2.1. SARS-CoV-2 ORF3a interacting human proteins

To find out influence of SARS-CoV-2 ORF3a in human cells, we sought to obtain cellular proteins interacting with SARS-CoV-2 ORF3a viroporin from Human protein atlas and collected the information about SARS-CoV-2 interacting human proteins.^51^ SARS-CoV-2 ORF3a viroporin interacts with eight human cellular proteins. To find out tissue specific effect of ORF3a viroporin, we started our analysis with these eight proteins, identified tissue specific proteins and miRNAs interacting with these eight proteins, analysed GO (Gene Ontology) term enrichment analysis to identify biological processes associated with these interacting networks.

### 2.2. Identification of hub genes and clustering of ORF3a interacting proteins network

ORF3a interacting interacting genes that are common in brain, heart and lung were used to construct protein protein interaction in STRING version 11.0.^52^ This network was further grown to obtain more interactions. Whole network was uploaded in Cytoscape version 3.9.0^53^ and hub genes for different network were found using Cytohubba plugin.^54^ In our study node scores of hub genes were calculated on the basis of Maximum Clique Centrality (MCC), bottleneck and EcCentricity algorithm separately. Protein protein interaction network of tissue specific common proteins in heart, lung and brain were further clustered into groups to find highly connected regions in the network using MCODE ^55^ application in Cytoscape. K-core cut off 2 was used with maximum depth of 100. K-core denotes the score deviance from the seed node’s score for exapanding the cluster. Maximum depth is the limit by which search distance from seed is set. Cluster modules network were separately analysed to find the biological processes regulated by seed proteins in clusters. BiNGO (Biological Networks Gene Ontology)^56^ tool in Cytoscape was used for assessing statistically over represented biological processes regulated by these clusters with significance value 0.05 as cut off. Hypergeometric test was used as statistical test and Benjamini and Hochberg false discovery rate (FDR) correction was used for multiple testing correction.

### 2.3. Tissue specific protein interaction network construction

Tissue specific interacting protein partners of these eight proteins have been isolated from IID (Integrated Interactions Database) (http://ophid.utoronto.ca/iid) based on two or more experimental evidences (studies or bioassays).^57^ The experiments in which these interactions were established are provided in supplementary material along with PubMed IDs. Tissue specific common and unique proteins (provided in supplementary materials) have been identified through Venn diagram analysis using http://genevenn.sourceforge.net/. Tissue specific interacting proteins have been used for further analysis. Gene ontology (GO) enrichment analysis was performed using the R package clusterProfiler.^58^ Benjamini and Hochberg test had been used for pAdjustMethod (for the FDR correction). Enrichments with p value ≤0.01 had been considered as significant. Enrichment for the common pathways, q value ≤ 0.01 was considered as significant.

### 2.4. Tissue specific miRNA network construction

ORF3a may regulate the activities of a diverse array of miRNAs. miRNAs are well known influencers of gene expression. Using miRNet (https://www.mirnet.ca)^59^ we have identified and constructed tissue specific interaction network of ORF3a interacting 8 proteins with miRNAs. Based on availability of data in miRNet, we only constructed miRNA networks for brain and lung. Degree of a node has been calculated by the number of other nodes it connected, and betweenness centrality signifies the bridging role of a node in a network. We have used Gene Ontology database for functional enrichment analysis of miRNA network and identified significant (p<0.05) biological processes associated with the miRNA networks. Enrichment analysis was based on the Hypergeometric tests after adjustment for FDR.

Likewise, as SARS-CoV-2 influences miRNAs, we wanted to study the miRNA-Protein network involving these miRNAs and SARS-CoV-2 influenced proteins interacting with SARS-CoV-2 ORF3a binding proteins. Common miRNAs in the reported list of SARS-Co-V2 influenced circulating miRNAs^60^ and the miRNAs that can target brain expressing interacting partners of SARS-CoV-2 ORF3a interacting partners were taken for analysis. Proteins taken for analysis were SARS-CoV-2 influenced SARS-CoV-2 ORF3a interacting proteins expressed in either brain or lung.

### 2.5. Delineating Tissue specific effect of ORF3a from RNA-seq data

Based on our above analysis, we sought to re-analyse published RNA-seq data to further delineate tissue specific effect of ORF3a. So, we chose three datasets including brain (Choroid Plexus Organoids study),^61^ lung (COVID-19 infected lung tissue compared to healthy subject),^62^ and heart-cardiomyocytes based SARS-CoV-2 transcriptomics study.^63^ Differentially expressed genes with P value ≤ 0.05 had been considered for further analysis. From these datasets, we selectively identified significant differentially expressed genes encoding proteins that directly or indirectly interact with SARS-CoV-2 ORF3a (here we considered the protein list depicted in table 1 as SARS-CoV-2 ORF3a interacting protein partners). Then we performed gene ontology enrichment analyses to find out relevant biological pathways in a similar way as depicted earlier. Gene ontology (GO) enrichment analysis had been performed in the similar way maintained earlier. Enrichments with p value ≤ 0.01 had been considered as significant.

**Table 1:** ORF3a directly interacting proteins.

|  | User ID | Ensembl Gene ID | Symbol | Gene Type | Species | Chr | Position (Mbp) |
| --- | --- | --- | --- | --- | --- | --- | --- |
| 1 | CLCC1 | ENSG00000121940 | CLCC1 | protein_coding | Human | 1 | 108.9274 |
| 2 | ARL6IP6 | ENSG00000177917 | ARL6IP6 | protein_coding | Human | 2 | 152.7179 |
| 3 | TRIM59 | ENSG00000213186 | TRIM59 | protein_coding | Human | 3 | 160.4324 |
| 4 | VPS11 | ENSG00000160695 | VPS11 | protein_coding | Human | 11 | 119.0678 |
| 5 | ALG5 | ENSG00000120697 | ALG5 | protein_coding | Human | 13 | 36.94974 |
| 6 | VPS39 | ENSG00000166887 | VPS39 | protein_coding | Human | 15 | 42.1587 |
| 7 | HMOX1 | ENSG00000100292 | HMOX1 | protein_coding | Human | 22 | 35.38036 |
| 8 | SUN2 | ENSG00000100242 | SUN2 | protein_coding | Human | 22 | 38.73473 |

## 3. **Results**

### 3.1. ORF3a interacting protein diversity varies in different tissues

SARS-CoV-2 affects various organs in different ways. To know this effect in brain, lung and heart, protein protein interaction studies were performed with proteins interacting with ORF3a interacting eight proteins. Proteins that were influenced by SARS-CoV-2 and those proteins that were expressed in brain, lung and heart were included in the analyses. These analyses from experiment-based tissue specific interaction network revealed 199, 197 and 175 interacting proteins in brain choroid plexus organoids, lung and heart respectively. Venn diagram analysis revealed 163 of these interacting proteins were common in all these three tissues. The analysis revealed ten unique interacting proteins (BLZF1, CDK5, ELOVL4, LRSAM1, NECAB2, SCARA3, SEC22A, TMEM17, UGT8, and ZDHHC22) in brain, seven unique interacting proteins (CHAT, CLDN4, CRB3, EVC2, PCDHB7, PDZK1IP1, and UNC93B1) in lung and only one unique interacting protein (EPN3) in heart (Supplementary data 1).

### 3.2. Analysis of SARS-CoV-2 influenced ORF3a interacting common proteins in brain, lung and heart predicted hub genes and clusters in the protein network

To find the prominent players in brain, lung and heart, we subjected the common SARS-CoV-2 influenced proteins for protein network identification. Protein protein networks constructed consisted of 168 nodes and 387 edges with average node degree of 4.61. Standard p-value ≤ 0.05 was used as cut off (Figure 1). Top 10 hub genes in the common proteins network identified by MCC algorithm were NOTCH1, HRAS, STAT3, ESR1, PECAM1, VIM, MKI67, PIK3CA, MAPK8 and VPS38 (Figure 2), whereas according to Bottleneck algorithm top 10 hub genes identified were NOTCH1, HRAS, PIK3CA, LAMP1, ESR1, MAPK8, KEAP1, GSN, ARHGDIA and YWHAB (Supplementary data 2). NOTCH1 and HRAS showed maximum significance among all hub genes. EcCenrtricity algorithm was also employed and top 10 hub genes identified were VDAC1, ESR1, MAPK8, BID, EZR, KEAP1, APP, BCAR1, ARHGEF1 and SQSTM1 that had the same rank (1.0) and score (0.333) (Supplementary data 2).

**Fig. 1.**
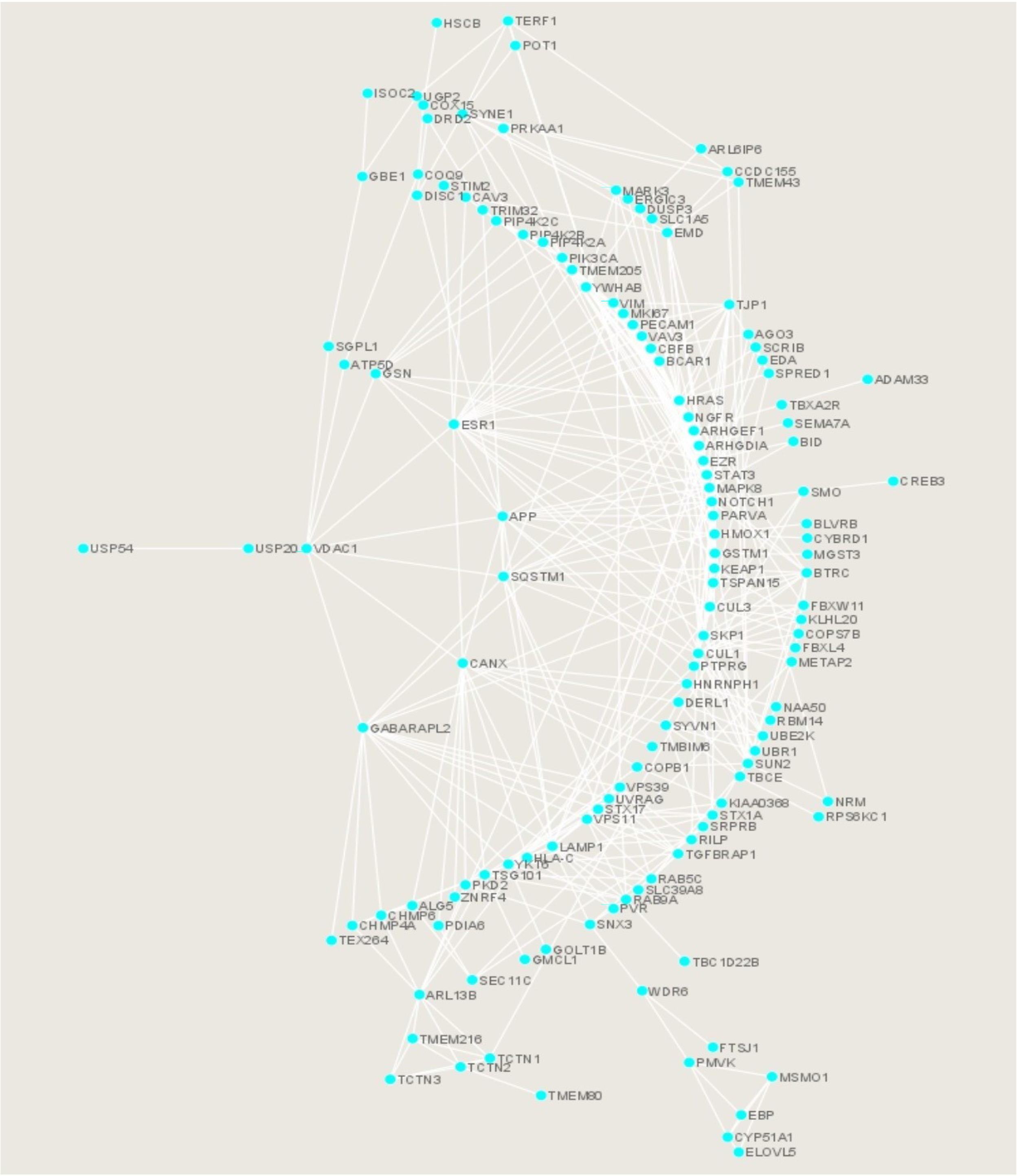
Network presentation of regulated common proteins expressed in the brain, lung and heart after SARS-CoV-2 infection. Visualization done in Cytoscape.

**Fig. 2.**
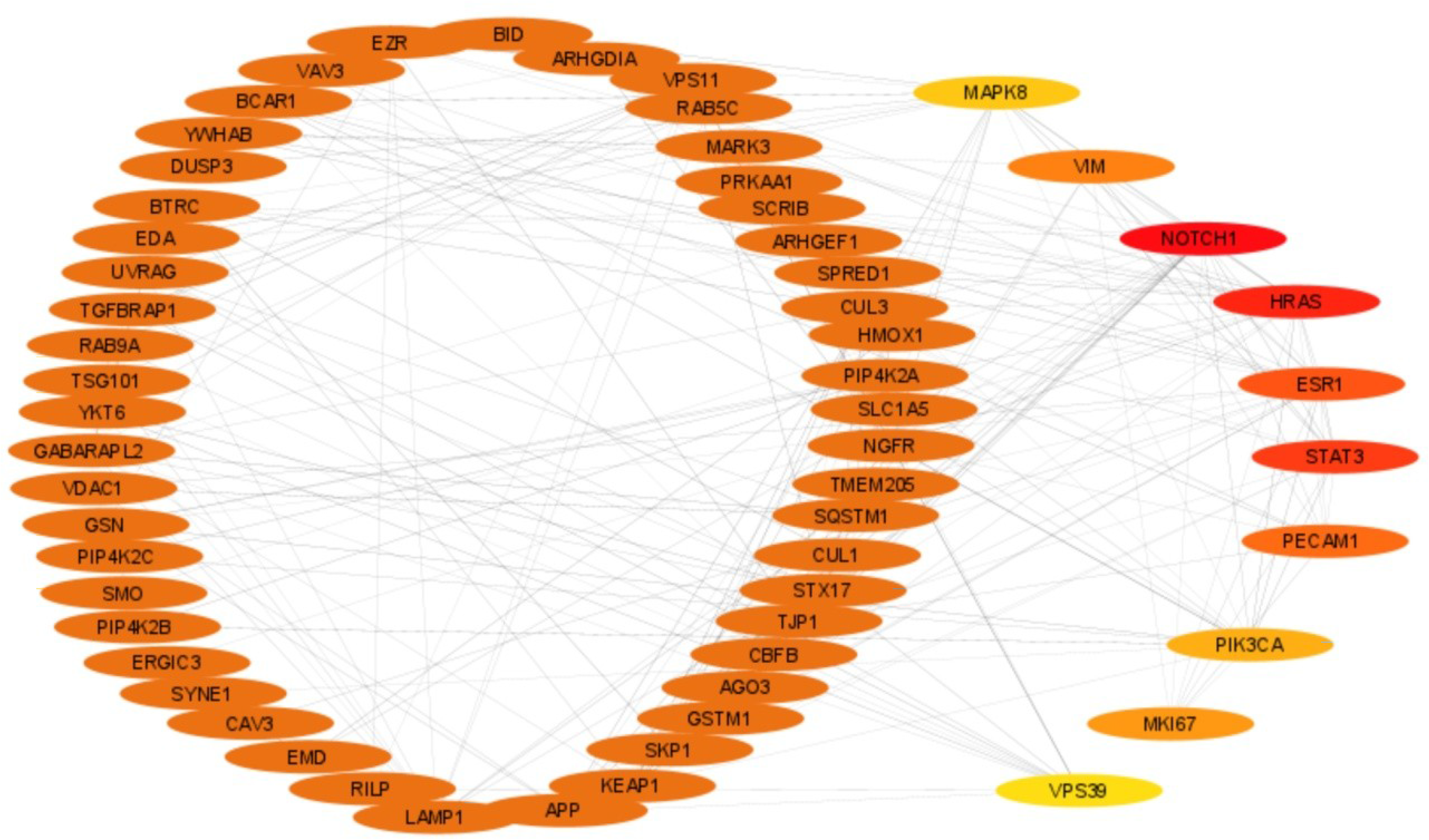
Top 10 hub genes of the common protein networks of regulated proteins after SARS-CoV-2 infection in the brain, lung and heart. Regulated proteins common in the brain, lung and heart were analysed by Cytohubba to find top 10 Hub genes (shown separately from the circularly arranged proteins). Node color from red to yellow represents higher to lower significance. NOTCH1, HRAS, STAT3 and ESR1 are highly significant major hub genes.

With the same data that we used to find the hub genes, MCODE was utilised to interpret highly connected top 5 clusters from the network (Figure 3 and supplementary data 3). For biological processes associated with every cluster, we used BiNGO. Cluster 1 had maximum score of 7.707 with two seed proteins, Proliferation marker protein Ki-67 (MKI67) and Ezrin (EZR) (Figure 4). Biological processes associated with it are regulation of cell death and apoptosis. Cluster 2 had maximum score of 6.182 with two seed proteins UV radiation resistance associated gene (UVRAG) and EZR (Figure 5). Maximum proteins in this complex regulate protein localization, transport and intracellular signal transduction. Cluster 3 (Figure 6), Cluster 4 (Figure 7) had score value of 3.600 and for Cluster 5 (Figure 8) it was 3.481. Seed proteins for them were Tectonic family member 2 (TCTN2) (Cluster 3), Methylsterol monooxygenase 1 (MSMO1) (Cluster 4) and again EZR for Cluster 5. Biological processes associated with Cluster 3 is protein transport, establishment of protein localization and other macromolecule localization. Cluster 4 is associated with regulation of cholesterol and sterol, steroid biosynthesis process, and lipid and alcohol biosynthesis. Biological process related to all five clusters are given in Supplementary data 4.

**Fig. 3.**
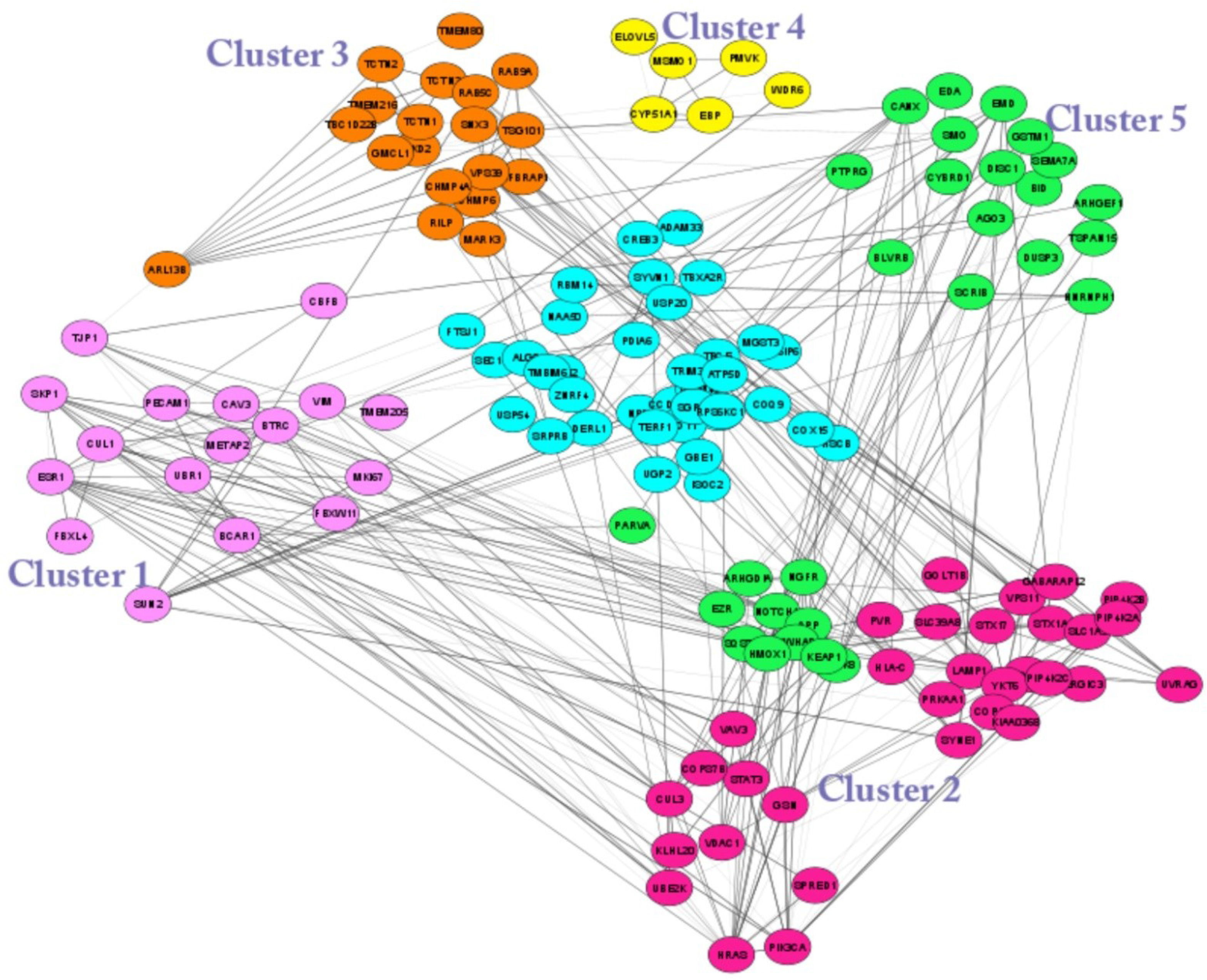
The MCODE analysis of protein protein interactome representing top five clusters. All five clusters are highly connected and each cluster is presented in different color.

**Fig. 4.**
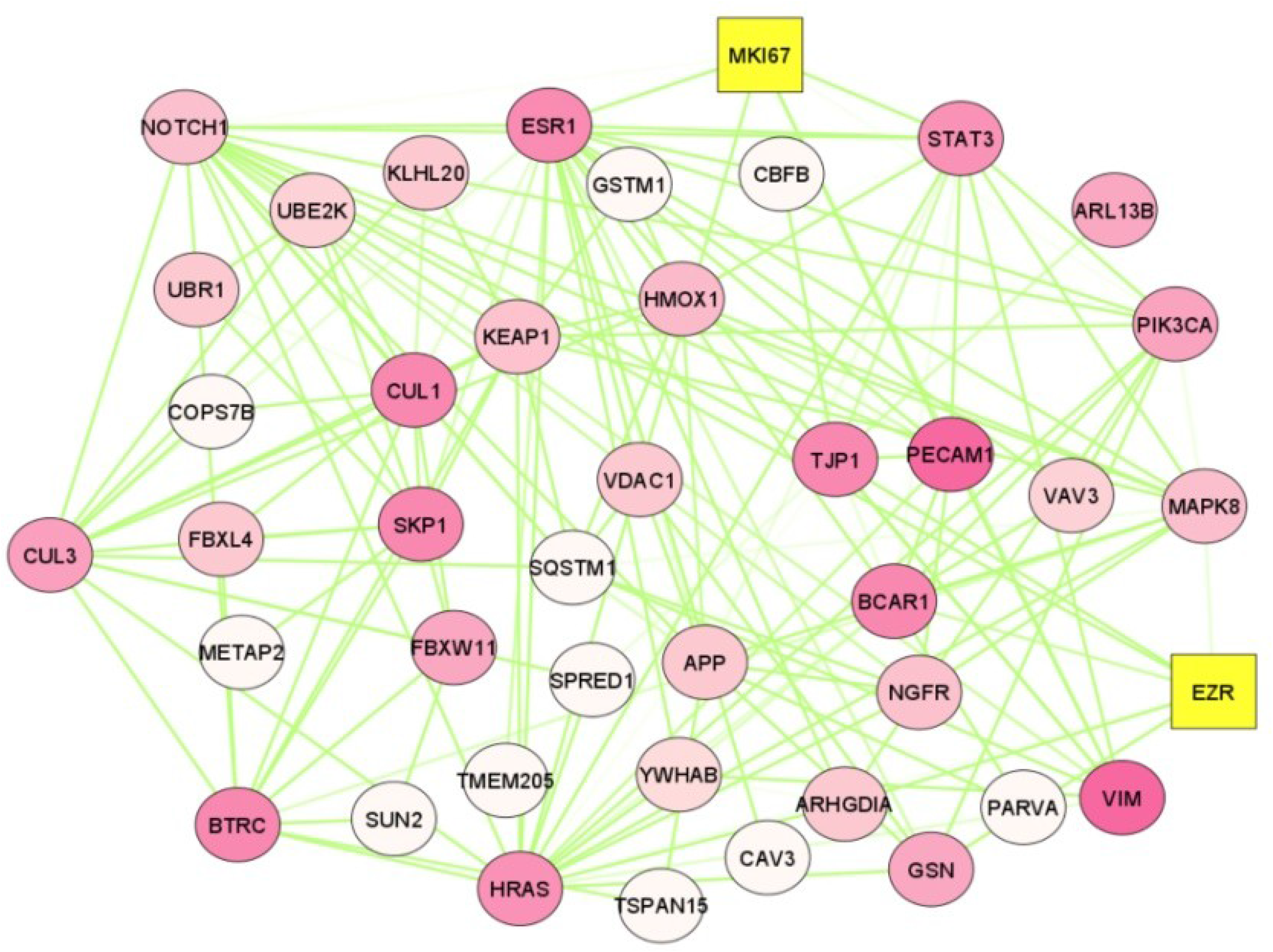
Cluster 1 (Score – 7.707 with 42 nodes and 158 edges). MKI67 and EZR are two seed proteins. This complex is responsible for regulating cellular processes like cell cycle, cell death, apoptosis and cellular metabolic process. Yellow coloured box denotes seed protein. Continuous mapping of node colour signifies maximum score with darkest shade (dark pink) to least significant with lightest shade.

**Fig. 5.**
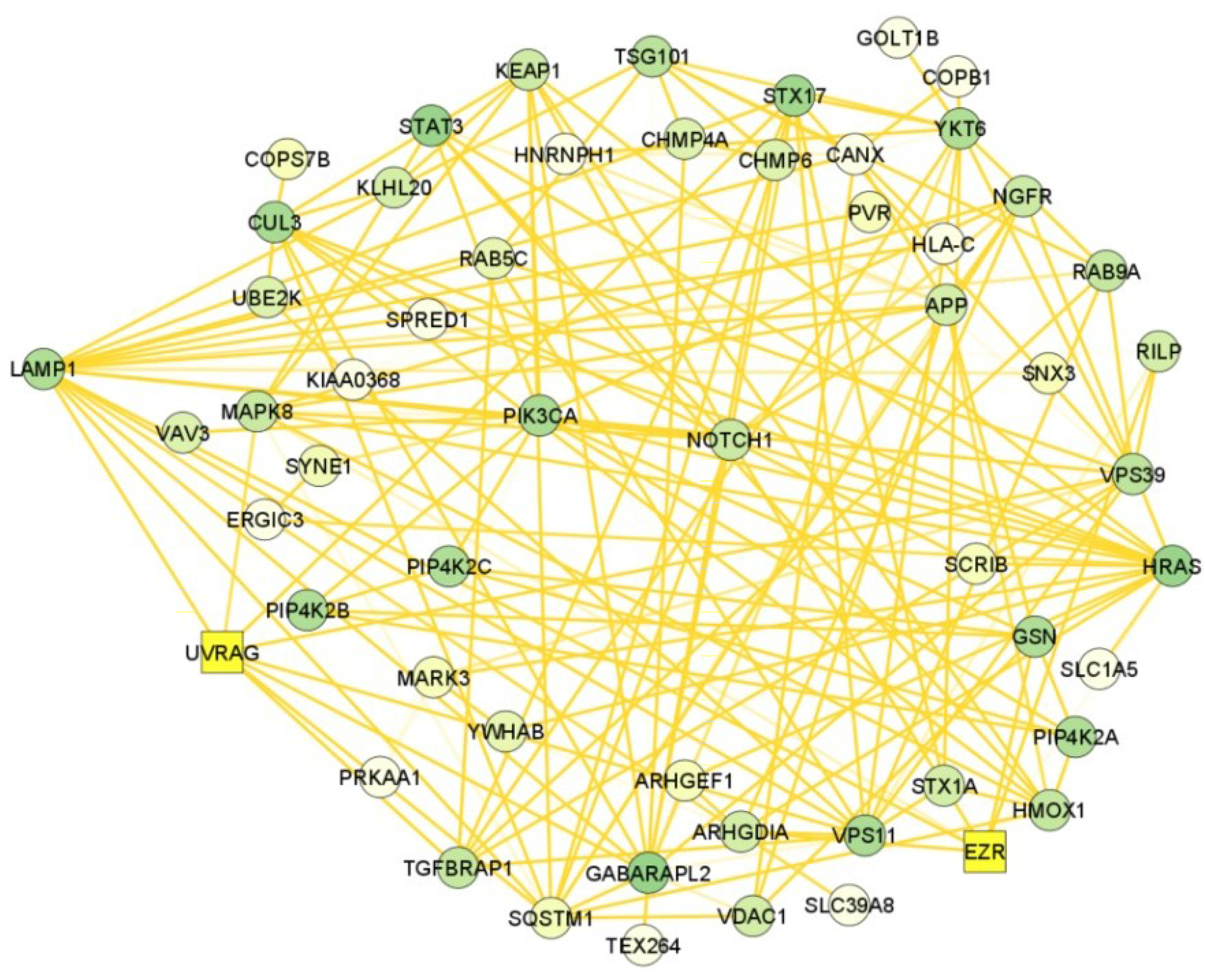
Cluster 2 (Score – 6.182 with 56 nodes and 170 edges). UVRAG and EZR are two important seed proteins in this cluster that are involved in cellular localization, protein localization, transport process as vesicle mediated transport and protein transport. Yellow coloured box denotes seed protein. Continuous mapping of node colour signifies maximum score with darkest shade (dark green) to least significant with lightest shade.

**Fig. 6.**
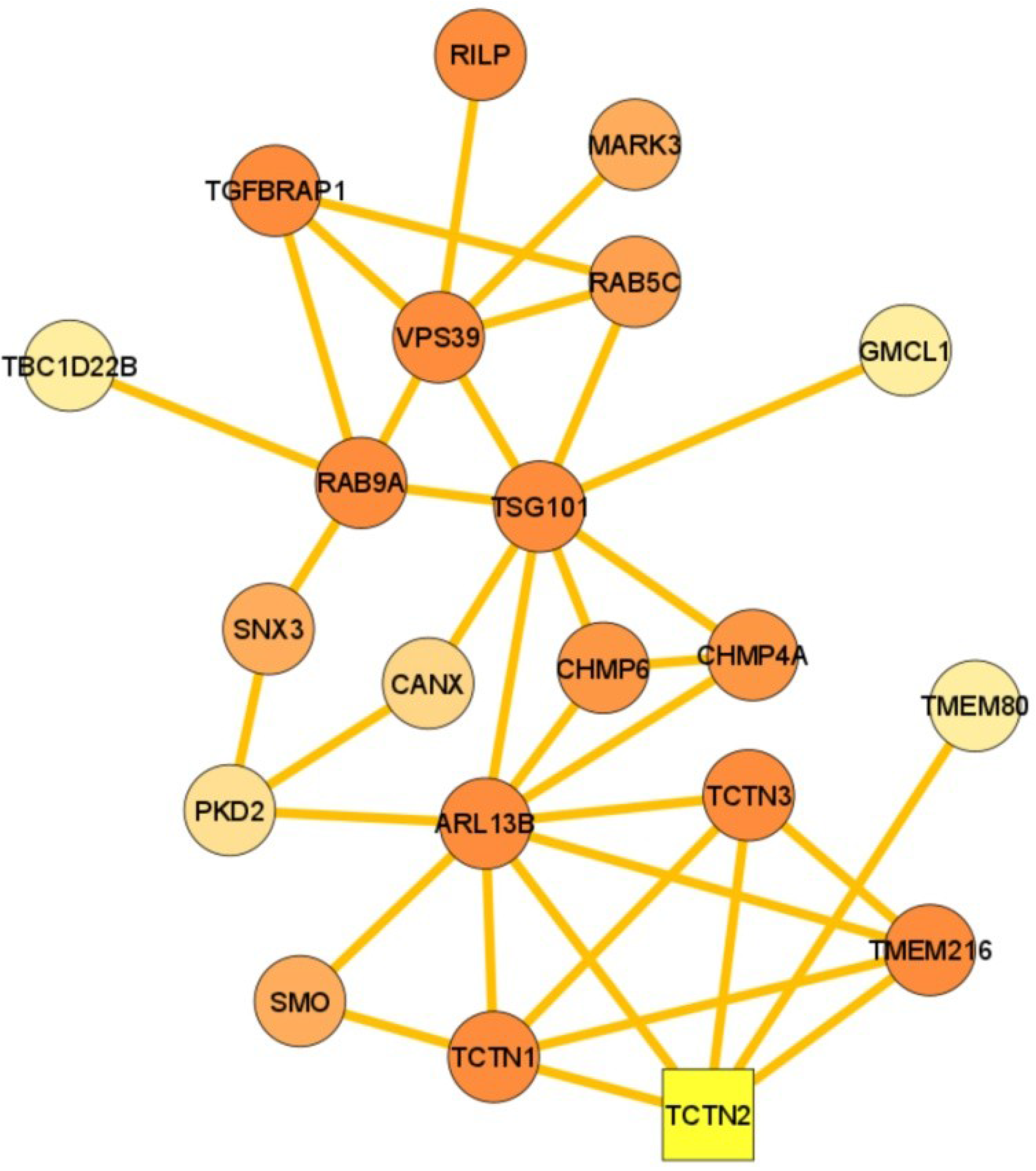
Cluster 3 (Score – 3.600 with 21 nodes and 36 edges). TCTN2 is acting as seed protein in this cluster regulating protein transport and localization processes. Yellow coloured box denotes seed protein. Continuous mapping of node colour signifies maximum score with darkest shade (dark orange) to least significant with lightest shade.

**Fig. 7.**
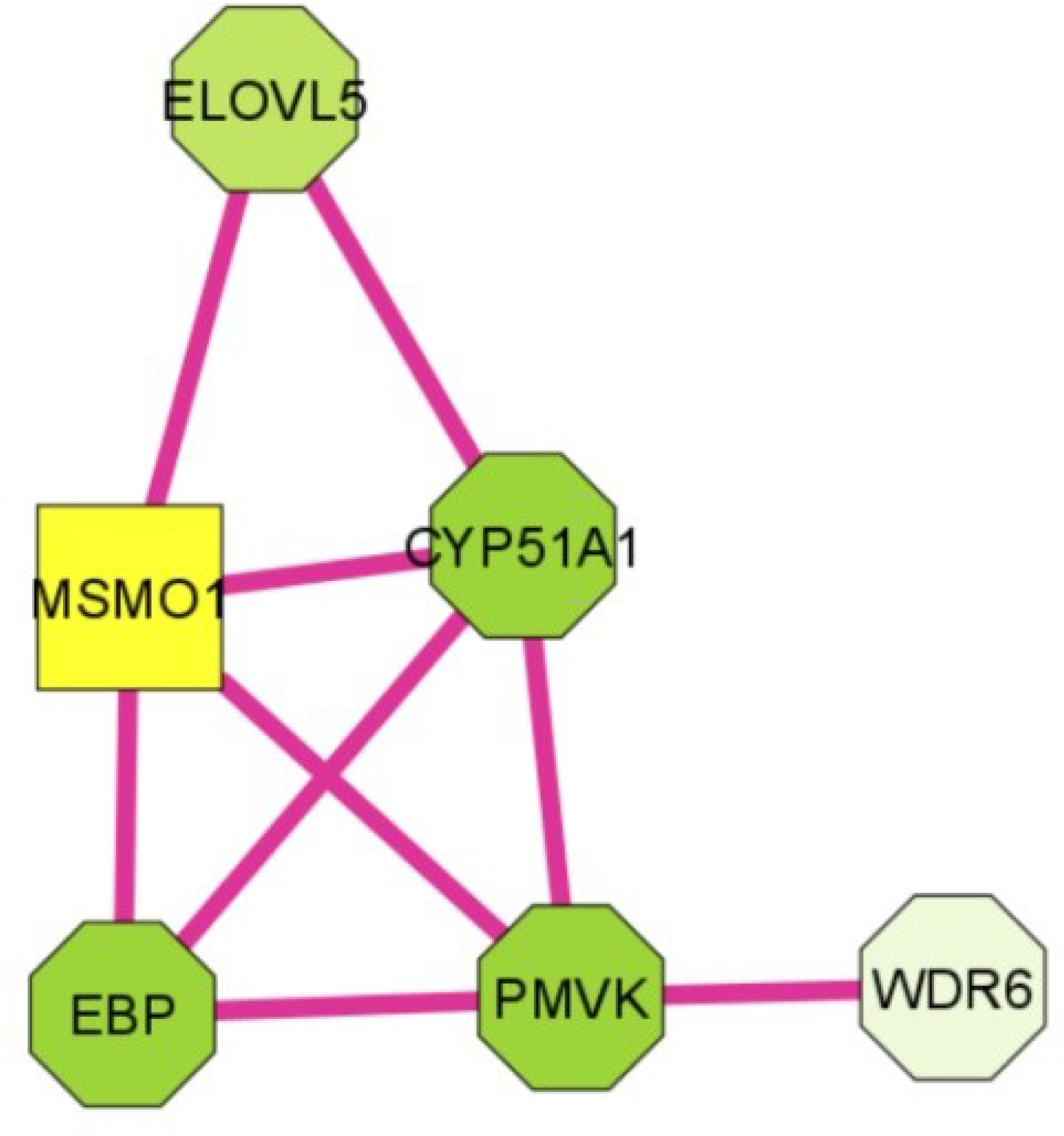
Cluster 4 (Score – 3.600 with 6 nodes and 9 edges). MSMO1 is the seed protein in this cluster regulating lipid and steroid biosynthesis along with alcohol, sterol, lipid and steroid metabolic process. Yellow coloured box denotes seed protein. Continuous mapping of node colour signifies maximum score with darkest shade (dark green) to least significant with lightest shade.

**Fig. 8.**
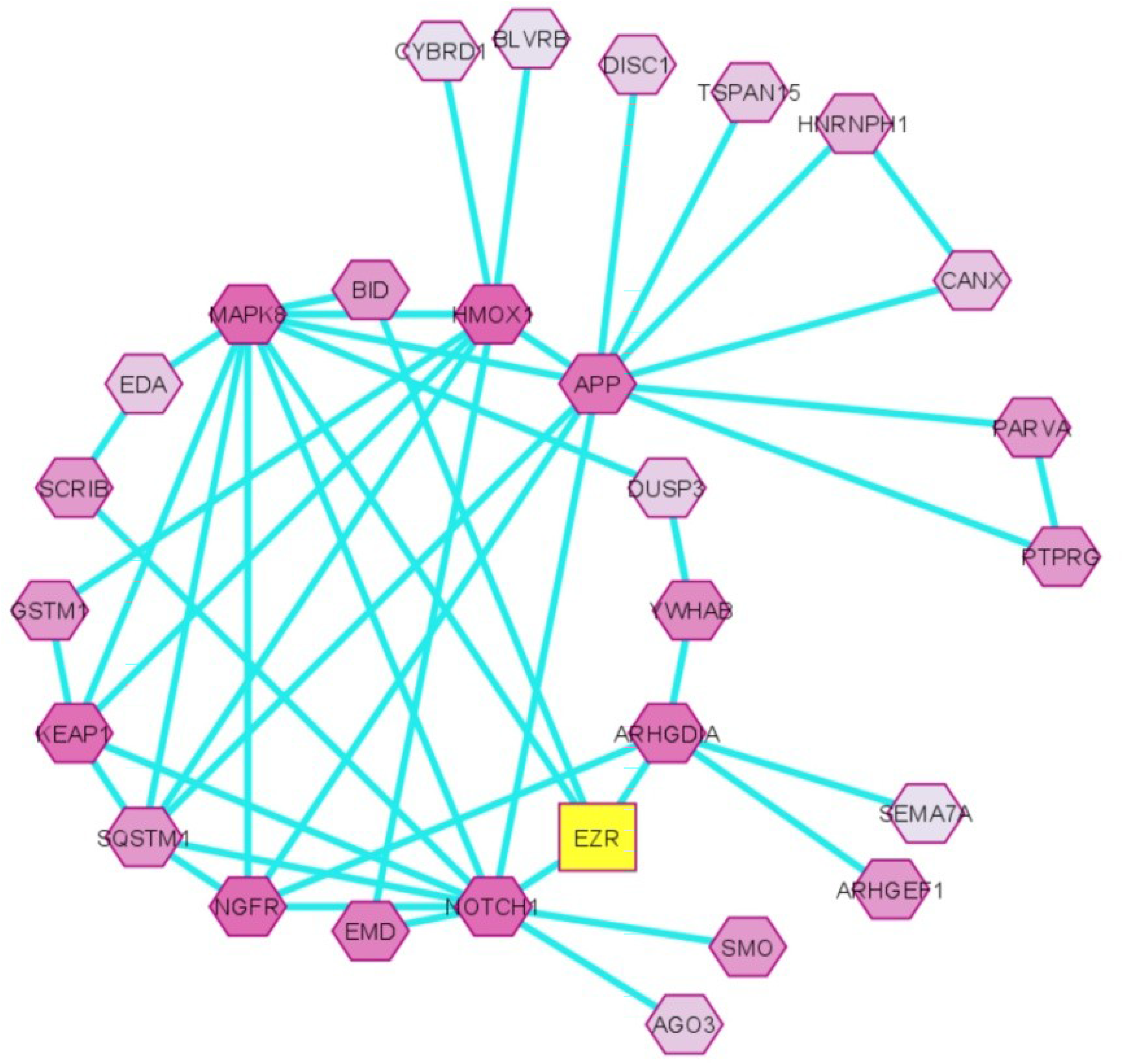
Cluster 5 (Score - 3.481 with 28 nodes and 47 edges). Seed protein of this cluster is EZR. Biological process annotation reveals cellular signalling and transduction and organ development processes. Yellow coloured box denotes seed protein. Continuous mapping of node colour signifies maximum score with darkest shade (dark violet) to least significant with lightest shade.

### 3.3. ORF3a interacting proteins can influence general and unique tissue specific biological processes

In general, ORF3a viroporin influences vesicular transport, cytoskeleton organization, protein kinase signalling, protein stability regulation and ubiquitination. Tissue specific interacting protein partners had been used to elucidate the related pathways that are probable and susceptible for COVID 19 infections in heart, brain and lung. Gene ontology enrichment analyses revealed no unique pathway in lung when compared to brain and heart. On the other hand, among these three major organs (heart, lung and brain), ORF3a mediated unique pathways were observed to be more in brain than in heart

(Figure 9). Common pathways influenced by ORF3a in all the three organs are shown in figure 10. In brain ORF3a influenced unique pathways include telencephalon and forebrain cell migration, intracellular transport, ERK1 and ERK2 cascade, controlling protein ubiquitination and localization, non-motile cilium assembly. Viral budding was one of the pathways observed to be affected only in brain (Figure 11) (Supplementary data 5 and 6). In heart ORF3a can also influence some unique biological pathways including the regulation of phosphatidyl inositol pathway, unfolded protein response in ER, positively regulate biotic stimulus and defence response. Among these, negative regulation of calcium ion transport by ORF3a interacting proteins can adversely affect cardiac function (Figure 12).

**Fig. 9.**
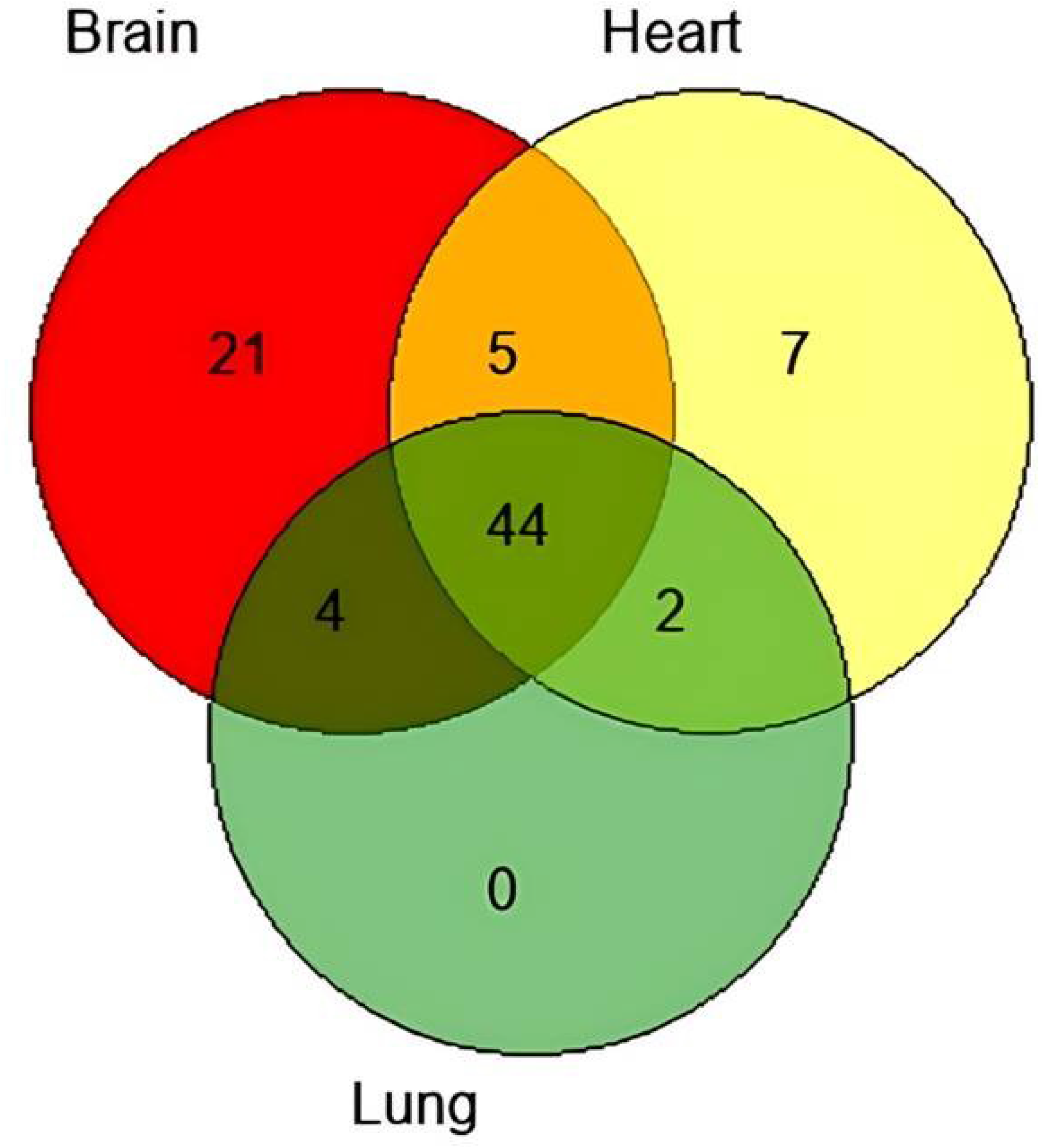
Venn diagram showing the number of biological pathways associated with ORF3a interacting proteins in heart, lung and brain. 44 pathways are common in heart, lung and brain. 21 unique pathways were observed in brain whereas in heart 7 were present. No unique pathway was observed in lung.

**Fig. 10.**
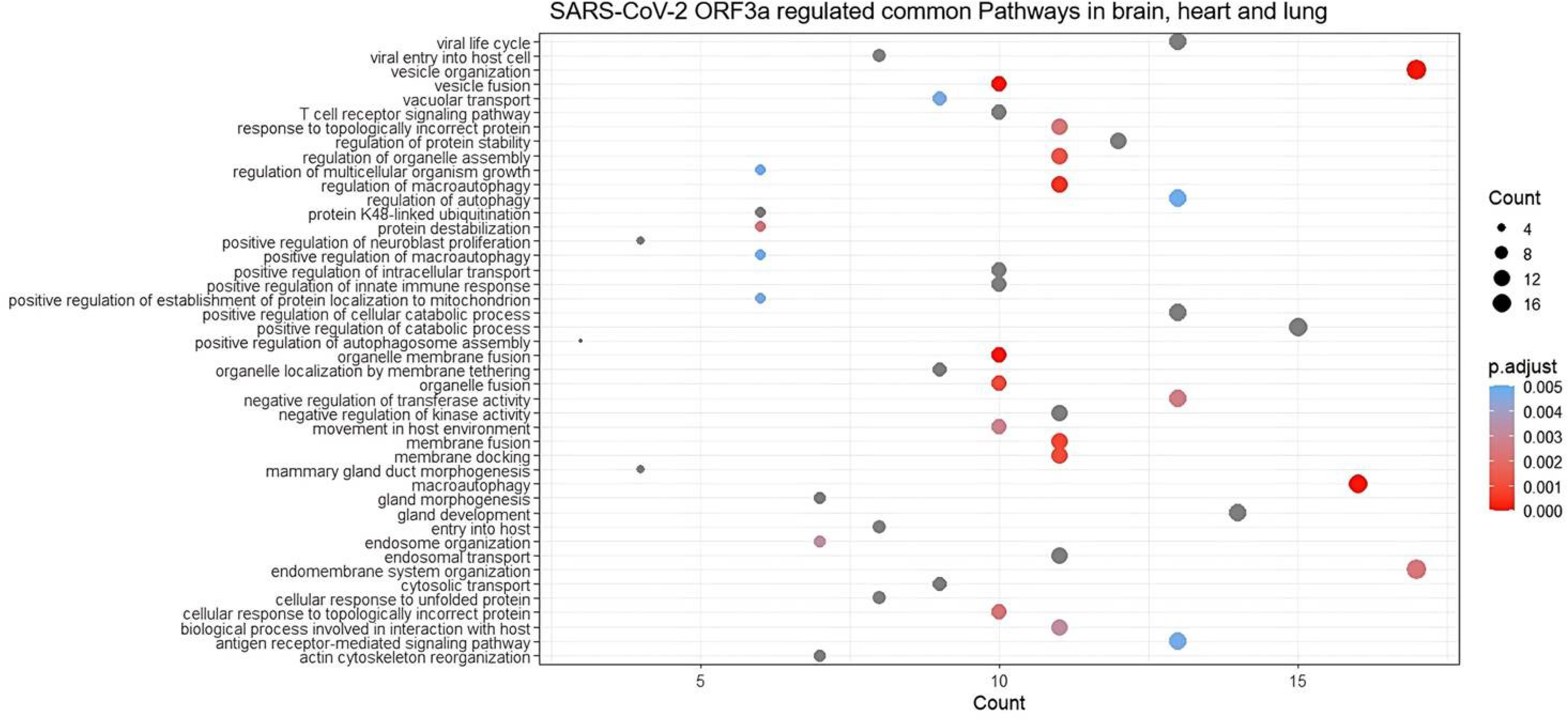
SARS-CoV-2 ORF3a regulated common pathways in lung, heart and brain. Dot size represents the gene ratio and dot colour depicts p.adjust value as in heat map.

**Fig. 11.**
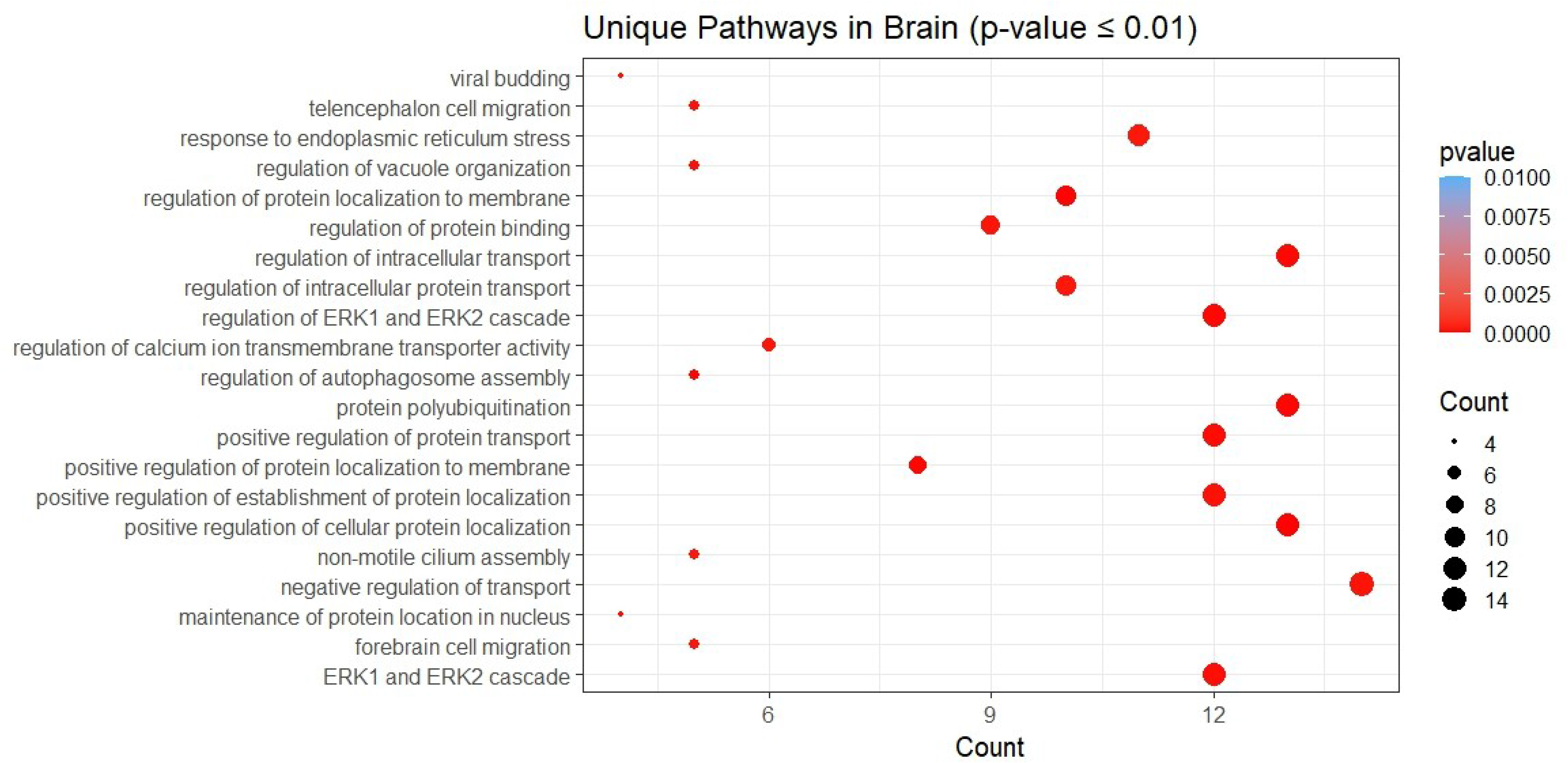
SARS-CoV-2 ORF3a regulated unique pathways in brain. Dot size represents the gene ratio and dot colour depicts P value as in heat map.

**Fig. 12.**
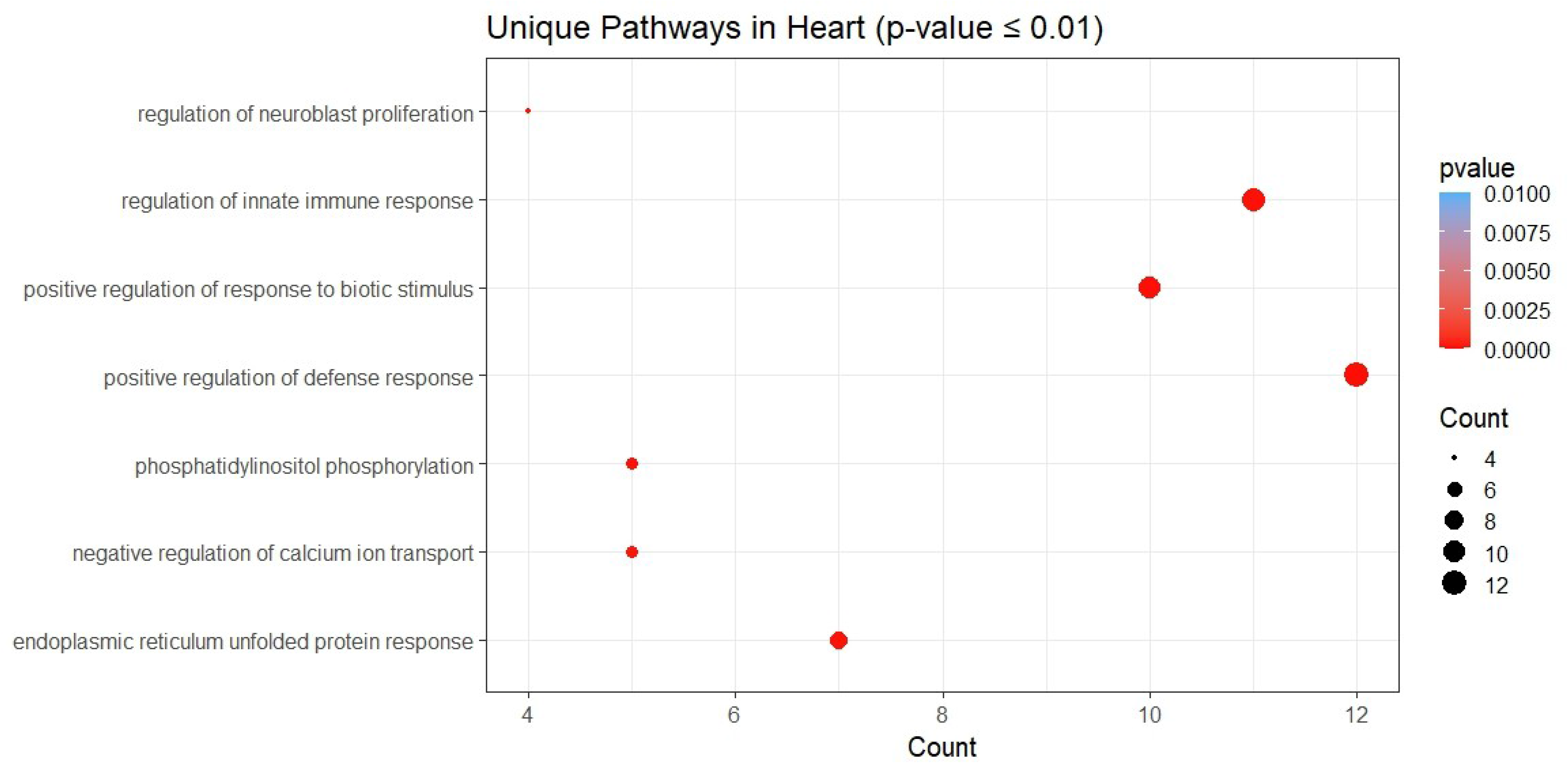
SARS-CoV-2 ORF3a regulated unique pathways in heart. Dot size represents the gene ratio and dot colour depicts P value as in heat map.

Next we wanted to analyse both upregulated and downregulated pathways based on SARS-CoV-2 influenced differentially expressed genes in a tissue specific manner. We delineated upregulated and downregulated genes to find out the pathways they influence. In brain choroid plexus organoid study, among upregulated genes influenced pathways prominent ones were negative regulation of phosphorylation, negative regulation of phosphate and phosphorus metabolic processes, and cellular response to peptide (Figure 13). Whereas, downregulated genes were observed to be associated with membrane docking, endosomal vesicular transport, and smoothened signalling pathway and these also deregulate neuronal patterning (Figure 14). In case of heart, upregulated genes were observed to be influencing immune response-regulating signalling pathway, phagocytosis, regulation of developmental growth, cellular carbohydrate metabolic processes, glycerophospholipid biosynthetic process, multicellular organism growth and regulatory processes, positive regulation of small molecule metabolic process, etc., (Figure 15). Downregulated proteins in heart are observed to influence positive regulation of intracellular protein transport, cholesterol biosynthetic process, endosome to lysosome transport, negative regulation of calcium ion transport, secondary alcohol biosynthetic process, selective autophagy, sterol biosynthetic process, etc. (Figure 16). Genes upregulated in lung were observed to influence biosynthetic processes of cholesterol, sterol and secondary alcohol (Figure 17). ERK1 and EFRK2 cascade and it’s negative regulation, macroautophagy, vacuolar transport, vesicle organisation, nucleus organisation, endosome organisation, membrane docking, protein K48-linked ubiquitination etc., were observed to be influenced by downregulated genes (Figure 18) (Supplementary data 7, 8).

**Fig. 13.**
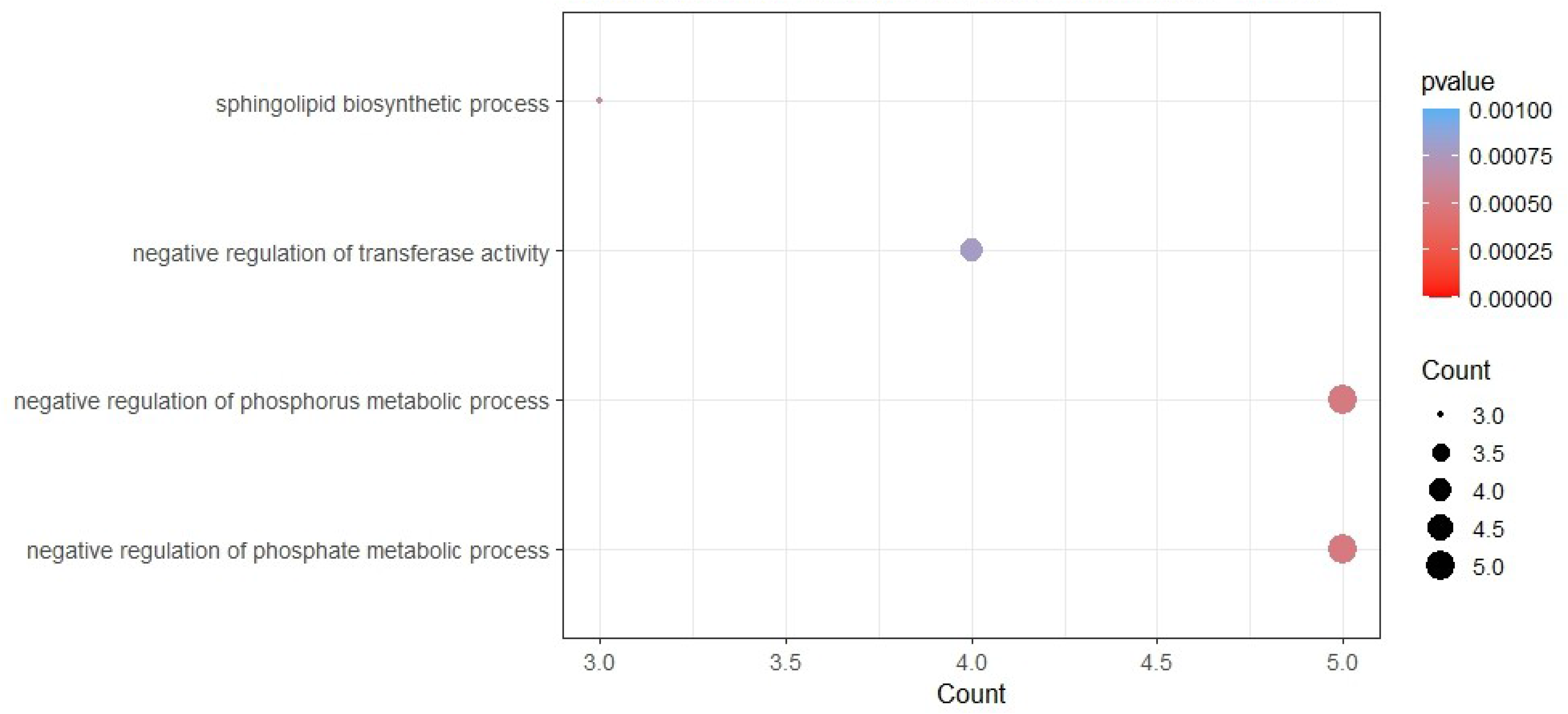
Dot plot of enriched Gene Ontology (GO) terms of differentially expressed upregulated genes in brain. Y-axis indicates the GO term and X-axis shows the count of genes per GO term. Colour gradient indicates p-value, using the Benjamini-Hochberg method.

**Fig. 14.**
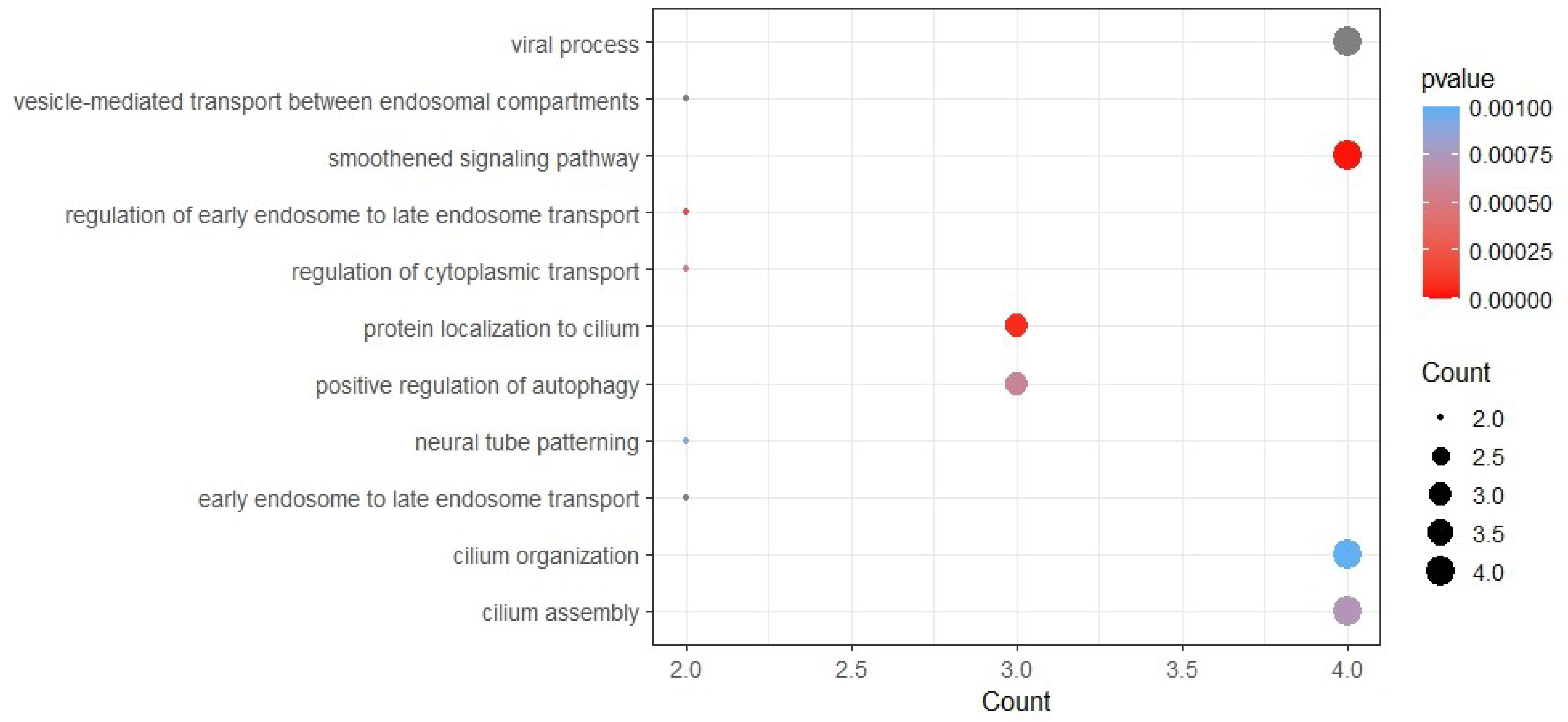
Dot plot of enriched Gene Ontology (GO) terms of differentially expressed downregulated genes in brain. Y-axis indicates the GO term and X-axis shows the count of genes per GO term. Colour gradient indicates p-value, using the Benjamini-Hochberg method.

**Fig. 15.**
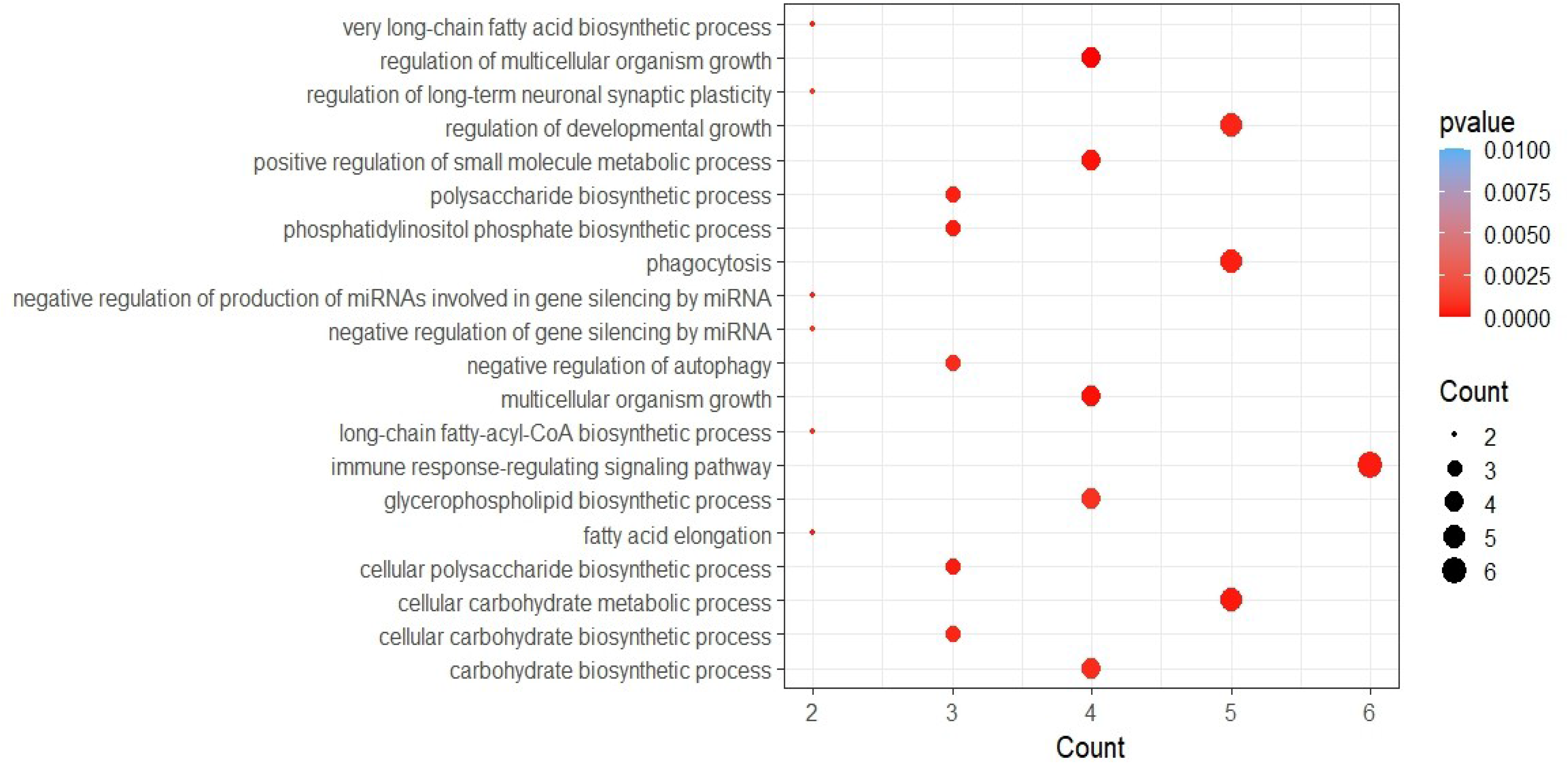
Dot plot of enriched Gene Ontology (GO) terms of differentially expressed upregulated genes in heart. Y-axis indicates the GO term and X-axis shows the count of genes per GO term. Colour gradient indicates p-value, using the Benjamini-Hochberg method.

**Fig. 16.**
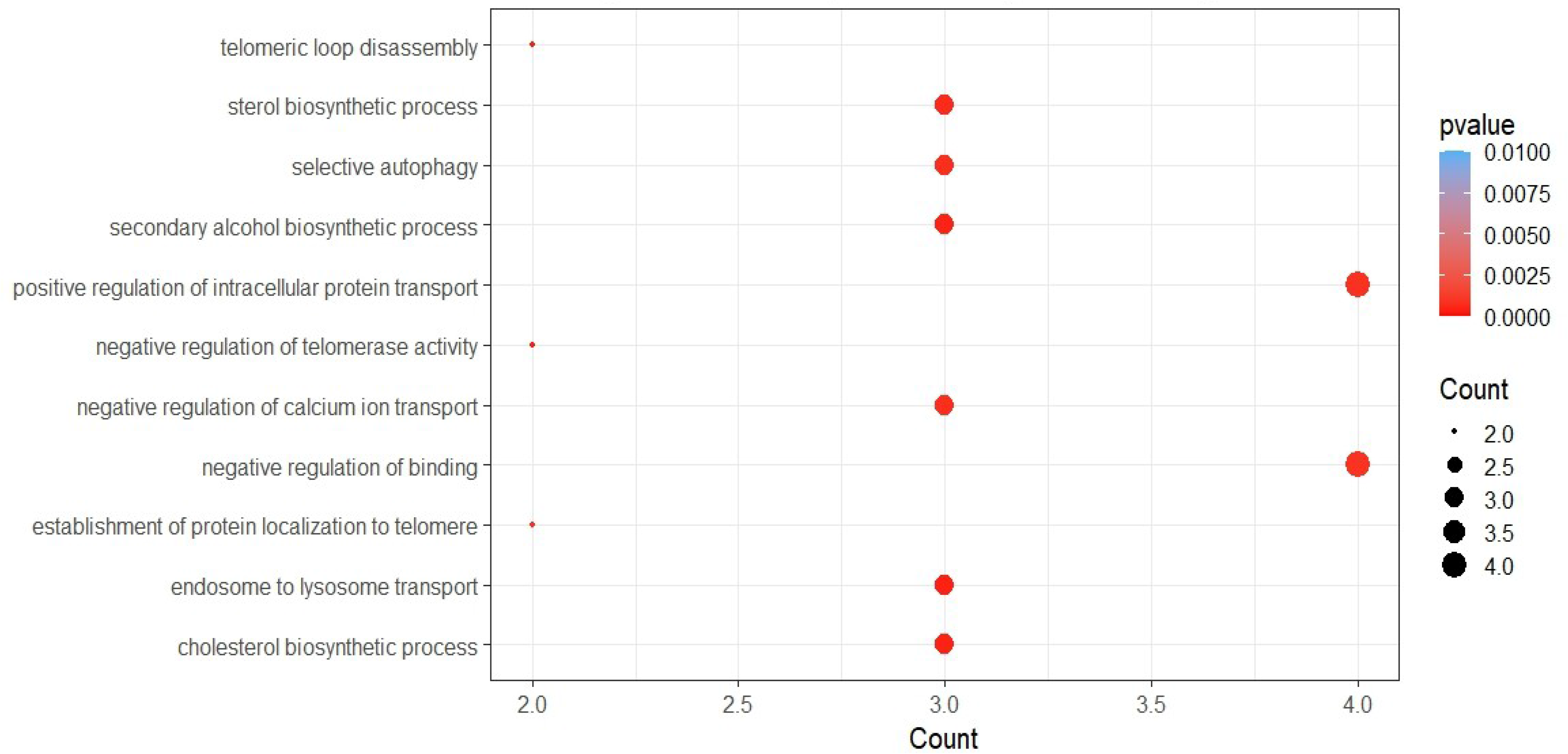
Dot plot of enriched Gene Ontology (GO) terms of differentially expressed downregulated genes in heart. Y-axis indicates the GO term and X-axis shows the count of genes per GO term. Colour gradient indicates p-value, using the Benjamini-Hochberg method.

**Fig. 17.**
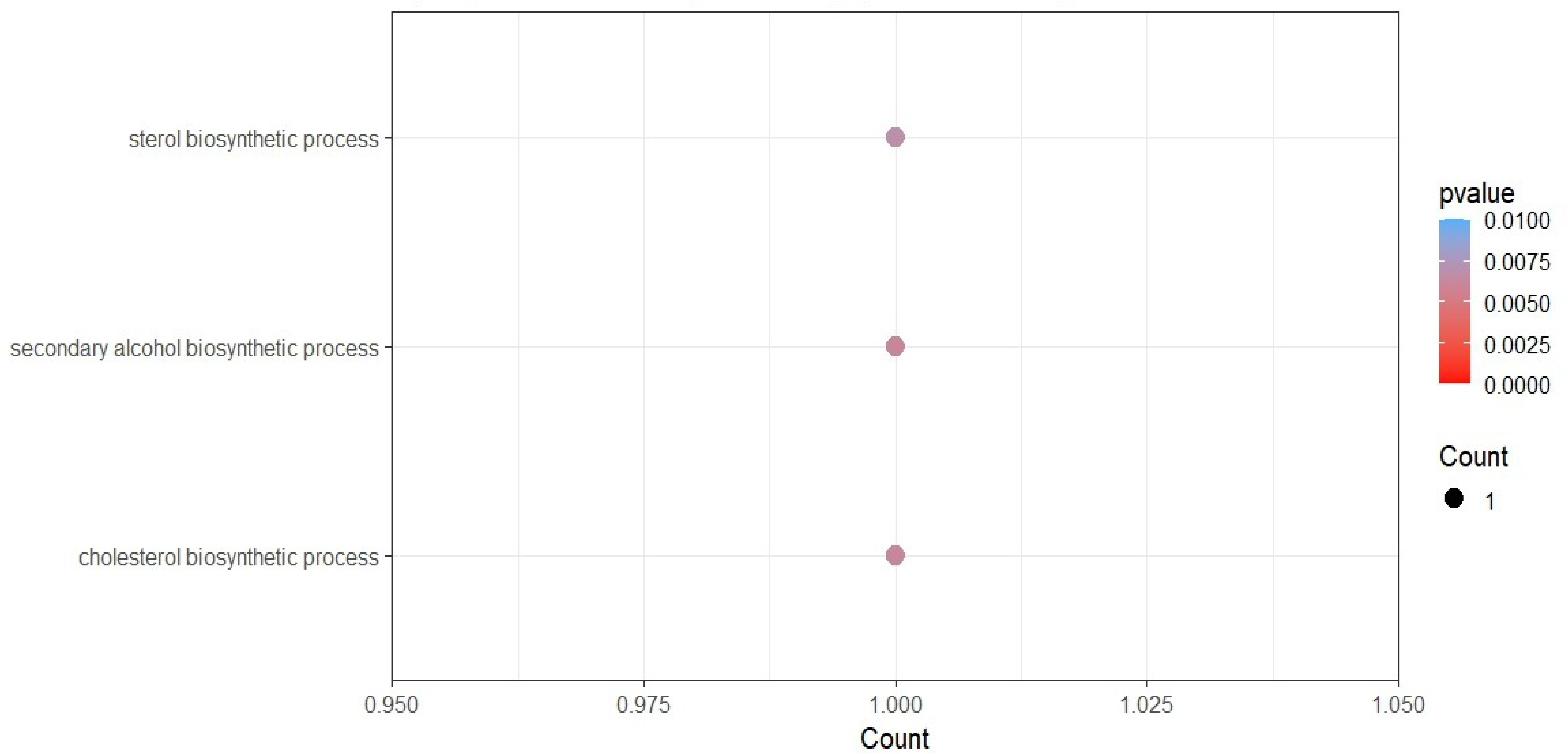
Dot plot of enriched Gene Ontology (GO) terms of differentially expressed upregulated genes in lung. Y-axis indicates the GO term and X-axis shows the count of genes per GO term. Colour gradient indicates p-value, using the Benjamini-Hochberg method.

**Fig. 18.**
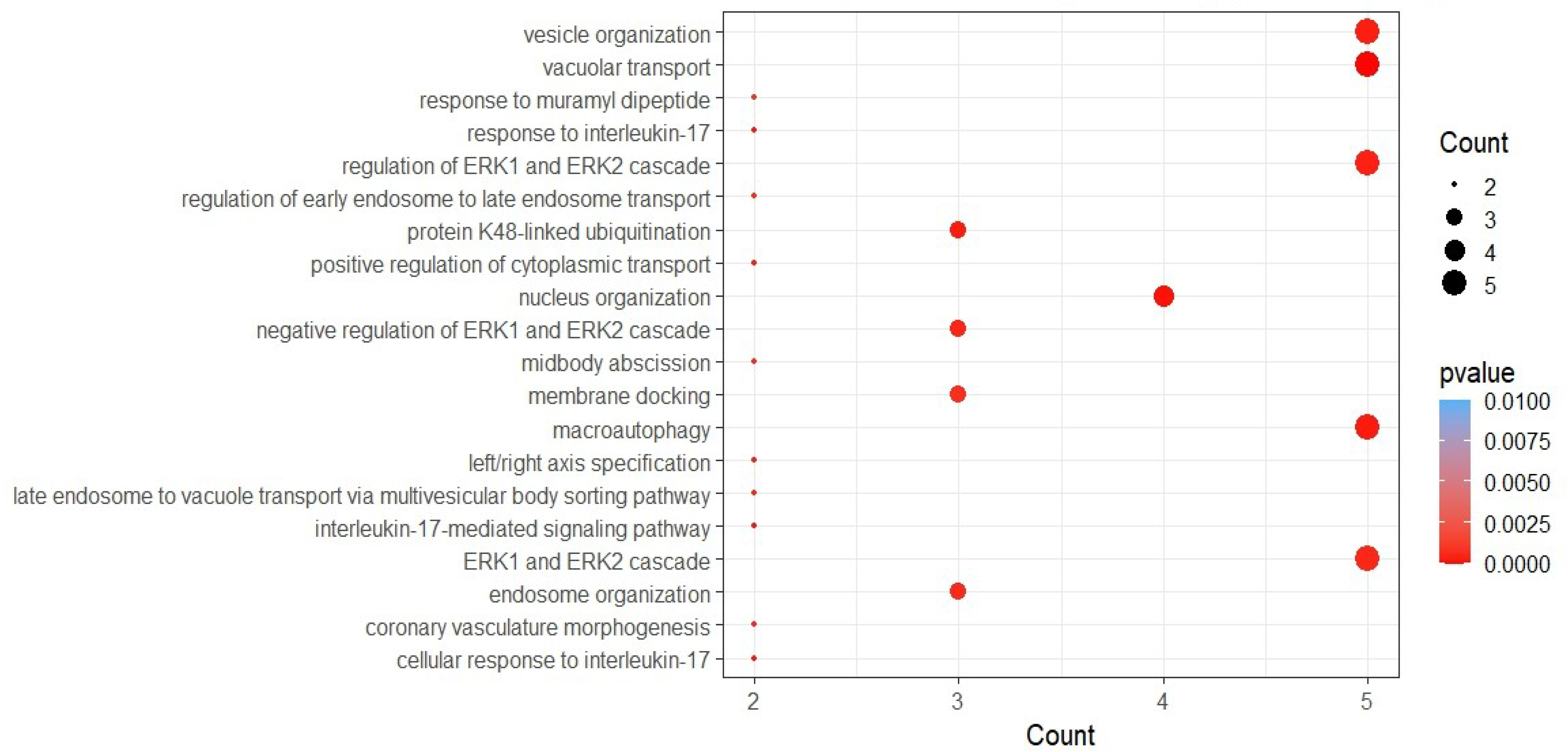
Dot plot of enriched Gene Ontology (GO) terms of differentially expressed downregulated genes in lung. Y-axis indicates the GO term and X-axis shows the count of genes per GO term. Colour gradient indicates p-value, using the Benjamini-Hochberg method.

### 3.4. ORF3a can influence biological processes of a tissue though miRNAs

As miRNAs influence protein levels, we wanted to find out miRNAs targeting SARS-CoV-2 ORF3a binding 8 proteins. From miRNet analysis we have identified 37 and 25 microRNAs interacting with ORF3a interacting eight proteins in brain (Figure 13) and in lung (Figure 14) respectively. In tissue specific miRNA interaction networks, we have identified two miRNAs with high degree and betweenness values (hsa-mir-1-3p with degree 7, betweenness 182.07, hsa-mir-124-3p with degree 6 and betweenness 122.92 and hsa-let-7b-5p with degree 5, betweenness 109.07) in brain. Similarly in lung specific miRNA interaction network we have identified only one miRNA with high degree and betweenness values (hsa-mir-1-3p with degree 7, betweenness 116.887). This indicates, that these miRNAs act as hub (for their high degree) and play a very important role in the interaction network (for their high betweenness). Apart from regulatory effect on cytoskeletal re-arrangement and protein localisation these miRNAs may influence diverse processes including NF-κB induced immune signalling cascades, cytokine biosynthesis, angiogenesis, cofactor catabolism, cell migration regulations, spindle fibre organization, DNA damage responses and endothelial cell proliferation as well as influences transcription factors binding with DNA. In brain, because of greater number of miRNAs interact with ORF3a interacting proteins, miRNA mediated pathways are more diverse. We observed three unique miRNA mediated biological pathways in brain. These were: protein N-linked glycosylation, cytokine metabolic process and negative regulation of sequence-specific DNA binding transcription factor activity (Supplementary data 9 – 14).

### 3.5. Tissue specific miRNA and protein interaction network of SARS-CoV-2 influenced miRNAs and proteins

To find out the effect of SARS-CoV-2 in host miRNA – ORF3a interacting protein network, we chose miRNAs that are reported to be regulated by SARS-CoV-2 with the following criteria. (i) miRNA should be present in the SARS-CoV-2 influenced circulating miRNA list,^60^ (ii) it should be expressed in our specific tissue of interest, and (iii) it should have a target in the expressed protein list of the tissue of interest that is influenced by SARS-CoV-2. To comply with the criteria, we first searched for miRNAs that can target the regulated proteins in miRNet for either brain or lung. Only those miRNAs that were present in the SARS-CoV-2 influenced circulating miRNA list were alone taken for network analysis. SARS-CoV-2 influenced proteins that are interacting with SARS-CoV-2 ORF3a interacting 8 proteins expressed in either brain or lung were taken for analysis to find out the miRNA-protein network in the respective tissues. In brain we found 4 miRNAs and they were hsa-let-7a-5p, hsa-let-7e-5p, hsa-miR-31-5p, and hsa-miR-651-5p (written in the order or decreasing degree). Among the proteins, WD repeat-containing protein 6 (WDR6) and signal transducer and activator of transcription 3 (STAT3) were targetted by all the 4 miRNAs (Figure 21). In case of lung, we could find only one miRNA hsa-mir-142-3p that was targetting many proteins (Figure 22). Common miRNAs that were observed to be targeting Orf3a interacting 8 proteins and it’s interacting proteins influenced by SARS-Co-V2 were hsa-let-7a-5p (targeting ARL6IP6) and hsa-mir-31-5p (targeting ALG5) in brain and in lung it was hsa-mir-142-3p (targeting ARL6IP6). Among the top 5 biological processes that could be regulated in brain are intracellular protein transport, tube morphogenesis, viral reproductive process, cellular macromolecule catabolic process and regulation of cellular protein metabolic process (Supplementary data 15). In lung we could find only two biological processes – I- kappaB kinase/NF-kappaB cascade and regulation of I-kappaB kinase/NF-kappaB cascade – with high significance (P – value ≤ 0.01) (Supplementary data 16).

**Fig. 19.**
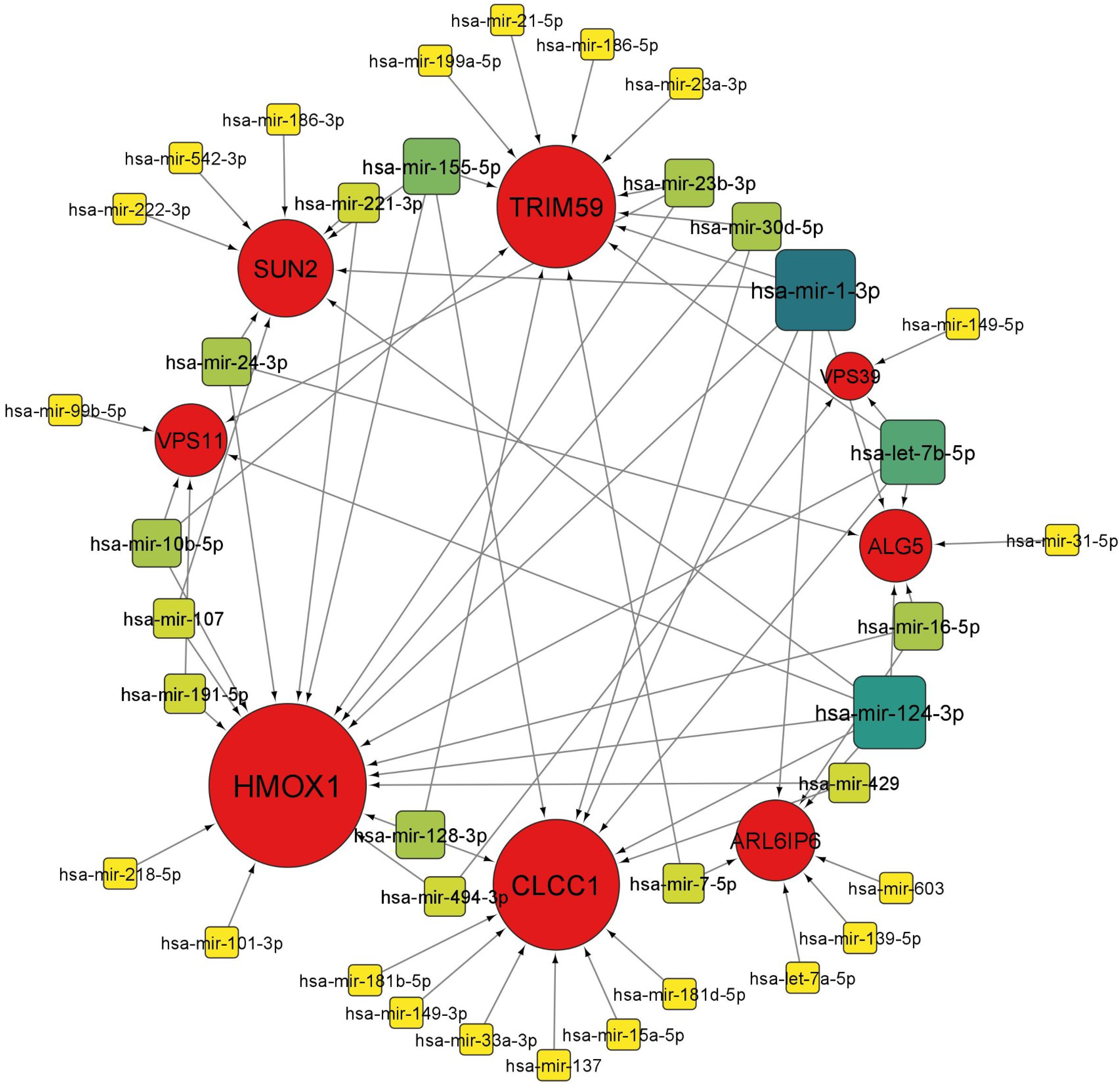
miRNA interaction network with SARS-CoV-2 ORF3a binding 8 proteins in brain. Circles represent ORF3a binding proteins and squares represent miRNAs. Increased size of nodes represent higher degree. For miRNA, darker shades of continuous mapping of node color represent higher significance (dark green to yellow).

**Fig. 20.**
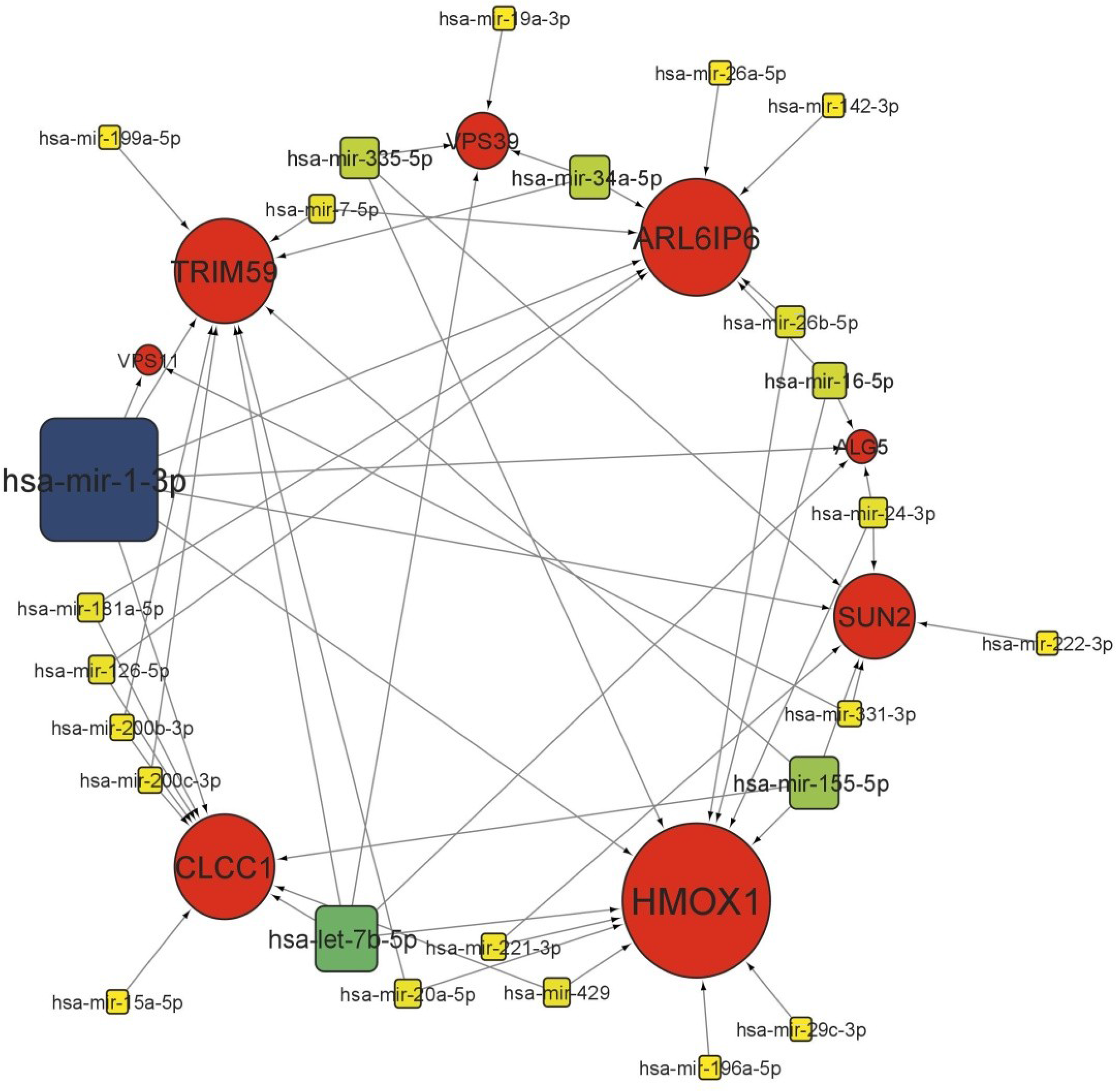
miRNA interaction network with SARS-CoV-2 ORF3a binding 8 proteins in lung. Circles represent ORF3a binding proteins and squares represent miRNAs. Increased size of nodes represent higher degree. For miRNA, darker shades of continuous mapping of node color represent higher significance (dark green to yellow).

**Fig. 21.**
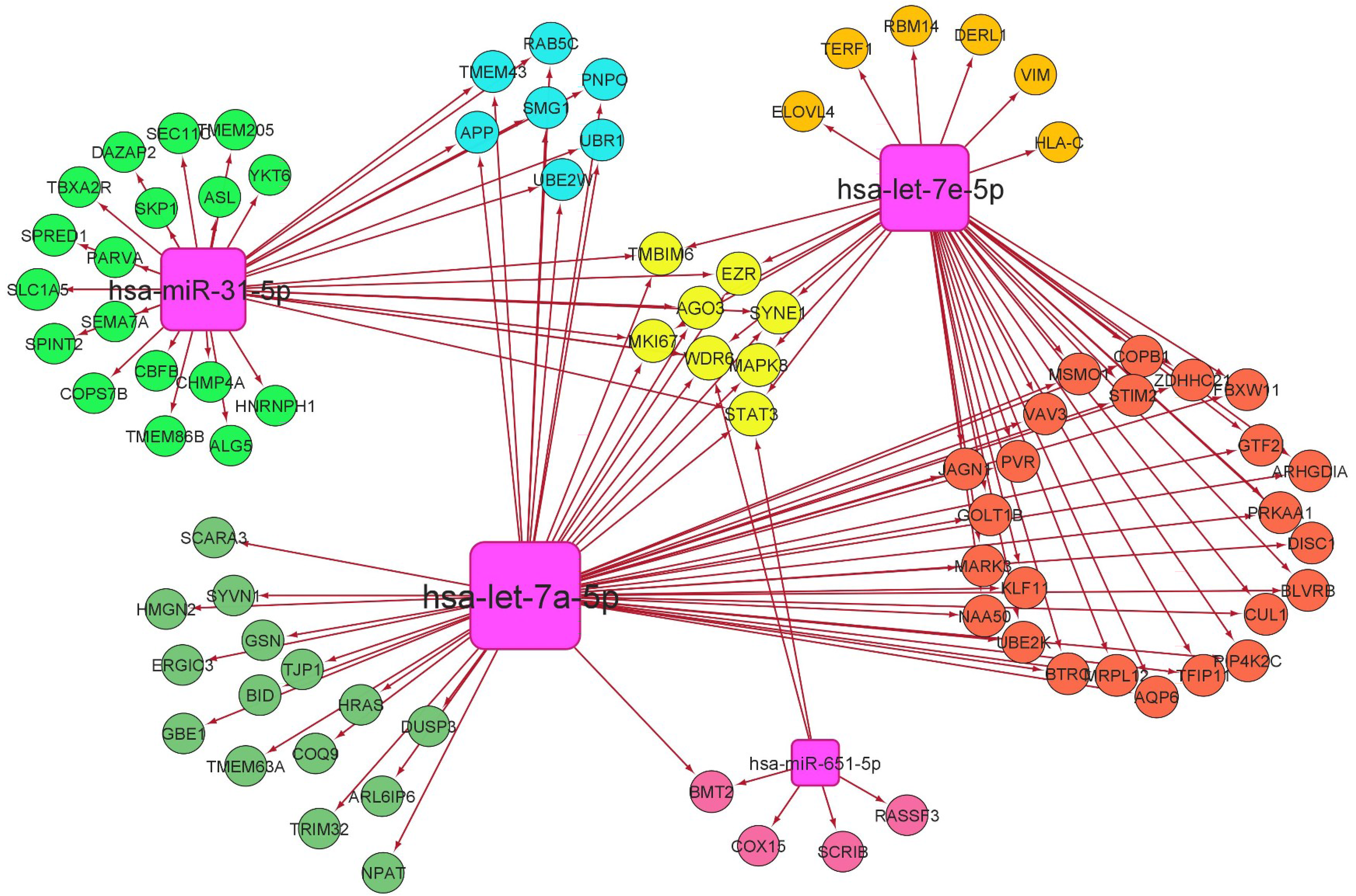
SARS-CoV-2 influenced miRNA – protein network in brain. Common miRNAs in the reported list of SARS-Co-V2 influenced circulating miRNAs^60^ and the miRNAs that can target brain expressing interacting partners of SARS-CoV-2 ORF3a interacting partners were taken for analysis. Proteins taken for analysis were SARS-CoV-2 influenced SARS-CoV-2 ORF3a interacting proteins expressed in brain. Nodes in circles are proteins and nodes in squares are miRNAs.

**Fig. 22.**
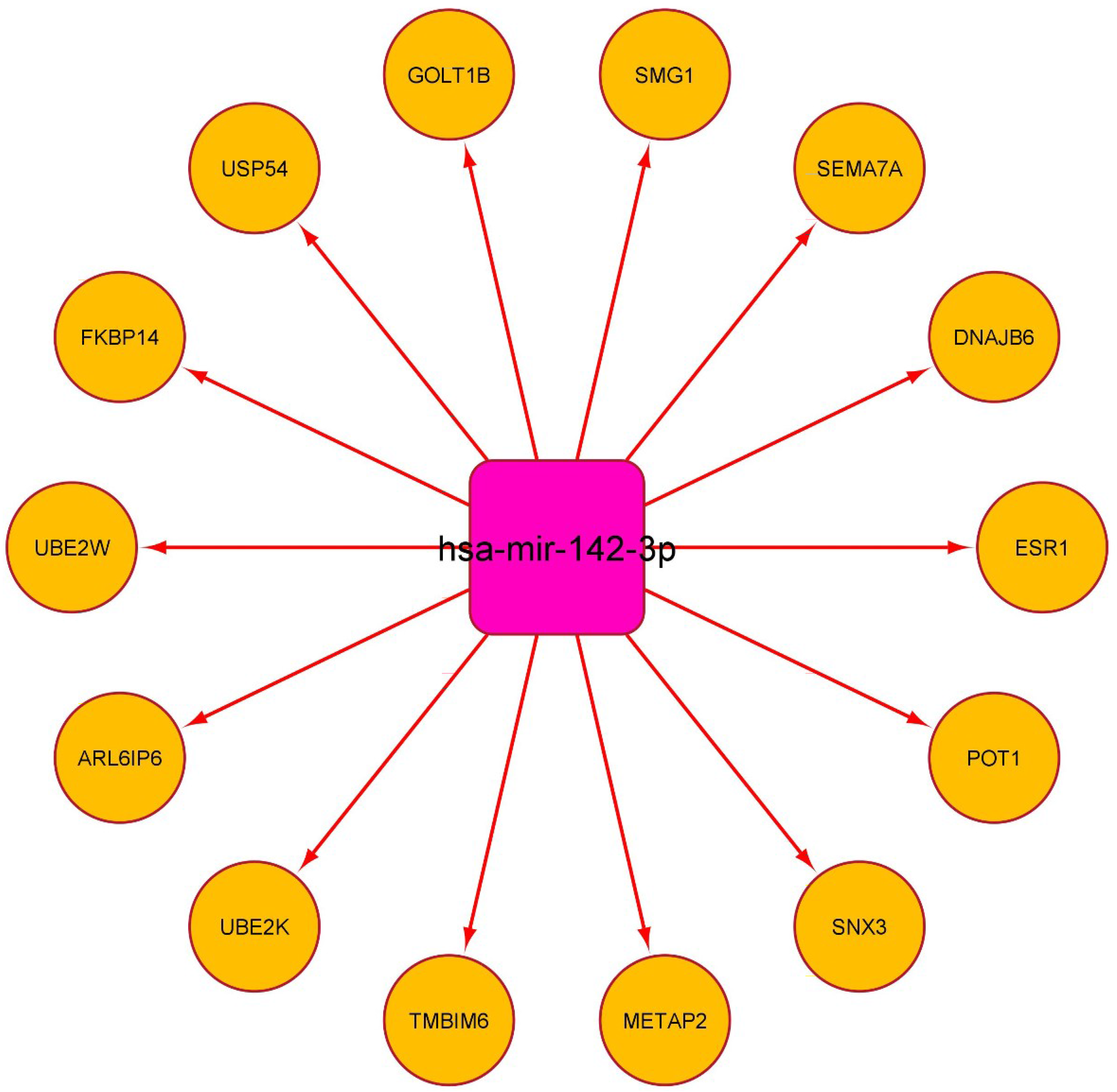
SARS-CoV-2 influenced miRNA – protein network in lung. Common miRNAs in the reported list of SARS-Co-V2 influenced circulating miRNAs^60^ and the miRNAs that can target lung expressing interacting partners of SARS-CoV-2 ORF3a interacting partners were taken for analysis. Proteins taken for analysis were SARS-CoV-2 influenced SARS-CoV-2 ORF3a interacting proteins expressed in lung. Nodes in circles are proteins and nodes in squares are miRNAs.

## 4. **Conclusions and Discussion**

Tissue specific complications upon SARS-CoV-2 infection had been delineated in several analyses. But the individual influence of SARS-CoV-2 accessory proteins are yet to be delineated in detail. Here we analysed human proteins interacting with ORF3a viroporin, extended the analyses by incorporating other interacting proteins interacting with ORF3a viroporin, and miRNAs interacting with ORF3a interacting proteins to get a global insight of ORF3a mediated responses in host cells. SARS-CoV-2 infection and expression of ORF3a in host cells can result in the initiation of general and tissue specific biological processes. We could find NOTCH1 and HRAS as hub genes in the common protein networks of regulated proteins after SARS-CoV-2 infection in the brain, lung and heart. NOTCH1 was also reported to be a major hub gene that plays multiple roles in SARS-CoV-2.^64,65^ HRAS observed to be involved in the SARS-CoV-2 immune response by peripheral blood mononuclear cells (PBMC).^66^ In our study, ORF3a interacting proteins were observed to be influencing pathways involved in the regulation of autophagy, macroautophagy and regulation of macroautophagy in the brain, heart and lung. It is reported that severe COVID-19 patients had significant impairment in antigen presentation with reduced expression of autophagy markers.^67,68^ Autophagy is one of the processes reported to be involved in MHC class I and II peptide presentation, thereby influencing antigen presentation.^69^ Induction of incomplete autophagy by ORF3a was shown to be through unfolded protein response.^70^ ORF3a and ORF7a were reported be co-localizing in late endosomes and preventing their acidification.^71^ Strong protein specific immunostaining of ORF3a was reported in plasma membrane and endosomes of SARS-CoV-2 infected Caco-2 cells.^72^ ORF3a was observed to be preventing the fusion between lysosomes and autophagosomes, whereas ORF7a reduces autophagosomal degradation by reducing the lysosome acidity.^71^ ORF3a inhibiting the fusion of autophagosomes with lysosomes is also reported.^49^ Other biological processes we observed to be influenced by OFR3a in heart, lung and brain are related to membrane and endomembrane systems. Biological processes involving vesicle organisation, membrane docking, membrane fusion, vesicle fusion, organelle membrane fusion and endosome organisation came in this category. SARS-CoV-2 ORF3a was reported to be localised in both membrane and cytosolic fractions^46^ like immunostaining that was shown in these compartments.^72^ It is also observed to be localised on late endosomes and unlike SARS-CoV ORF3a, directly interacts with HOPS component VPS39 leading to the prevention of HOPS interaction with autophagosome-localised STX17 culminating in the formation of autolysosome formation.^48^ In a later study, it had been depicted that systemic inflammation results the dysregulation of autophagy and neuroinflammation,^67^ as well as increased protein ubiquitination in SARS-CoV-2 infection.^73^ Our study adds the influence of ORF3a in K48- linked polyubiquitination as another common cellular events in SARS-CoV-2 infected in the brain, heart, and lung.

Proteomic study comparing ubiquitin modified proteome of SARS-CoV-2 infected cells speculates the modulation of K48-linked polyubiquitination to increase USP5 expression and type I IFN signal inhibition.^73^ In brain, effect of ORF3a can be severe as ORF3a interacting proteins influence cilia organisation and viral budding. These effects can initiate other tissue specific implications that can increase susceptibility to other diseases.

Another observation that we had in specific biological process in heart was negative regulation of calcium ion transport. Among COVID-19 patients treated with (298) and without (568) calcium channel blockers (CCBs), it was found that patients treated with CCBs had significant elevated rate of intubation.^74^ Another study among COVID-19 patients with a history of hypertension on dihydropyridine CCBs (70/245) and without CCB (170/245), there was a significant increase in the risk for intubation or death among those who were taking dihydropyridine CCBs.^75^ Another study showed that chronic CCB users were more likely to be hospitalized with COVID-19 in comparison with long term ACE (Angiotensin Converting Enzyme) inhibitors or ARBs (Angiotensin Receptor Blockers) using patients.^76^ Proteins involved in the biological process of negative regulation of calcium ion transport are Voltage-dependent anion-selective channel protein 1 (VDAC1), Transmembrane BAX inhibitor motiff containing protein 6 (TMBIM6) and Caveolin 3 (CAV3) where all three are down regulated. VDAC1 transports cations including Ca^2+^ ions in low conductance state.^77^ TMBIM6 was observed to be downregulated in SARS-CoV-2 infected cells^78^ which may prompt Ca^2+^ disorder in the cells.^78^ CAV3 was shown to regulate ion channels in caveolae of cardiac cells.^79^ L-type CCBs were shown to inhibit SARS-CoV-2 entry and infection in Vero E6 and Calu-3 cell culture.^80^ Repurposing these drugs could have deleterious effects evidenced by observed reports^74–76^ and our observation also seems to be back the observed reports. Though further in depth studies are necessary to investigate the role of ORF3a in this aspect, cumulative effect of downregulation of these three proteins could be playing an important role in the observed deleterious effect of CCBs in the context of SARS-CoV-2 infection. Our analyses included three datasets and even with that we were able to find out many tissue specific biological processes. Detailed transcriptomics analyses of large number of COVID-19 patients could unravel the tissue specific influence of ORF3a in the severity of COVID-19 infection in much more detail.

## 5. **Supplementary materials**

Supplementary Data 1: All interacting proteins

Supplementary Data 2: For Hub genes

Supplementary Data 3: PPI network with cluster parameters

Supplementary Data 4: Various clusters

Supplementary Data 5: ORF3a related all DEGs biological pathways

Supplementary Data 6: ORF3A related common and unique pathways

Supplementary Data 7: ORF3a DEGs biological pathways

Supplementary Data 8: ORF3a related all DEGs in heart lung brain

Supplementary Data 9: Brain mirnet_enrichment

Supplementary Data 10: Brain mirnet_mir_gene

Supplementary Data 11: Brain node_table_mirnet

Supplementary Data 12: Lung mirnet_enrichment

Supplementary Data 13: Lung mirnet_mir_gene

Supplementary Data 14: Lung node_table_mirnet

Supplementary Data 15: Brain_mirnet_enrichment_ORF3a int proteins

Supplementary Data 16: Lung_mirnet_enrichment_ORF3a int proteins

## Supporting information

Supplementary Data 1: All interacting proteins

Supplementary Data 2: For Hub genes

Supplementary Data 3: PPI network with cluster parameters

Supplementary Data 4: Various clusters

Supplementary Data 5: ORF3a related all DEGs biological pathways

Supplementary Data 6: ORF3A related common and unique pathways

Supplementary Data 7: ORF3a DEGs biological pathways

Supplementary Data 8: ORF3a related all DEGs in heart lung brain

Supplementary Data 9: Brain mirnet_enrichment

Supplementary Data 10: Brain mirnet_mir_gene

Supplementary Data 11: Brain node_table_mirnet

Supplementary Data 12: Lung mirnet_enrichment

Supplementary Data 13: Lung mirnet_mir_gene

Supplementary Data 14: Lung node_table_mirnet

Supplementary Data 15: Brain_mirnet_enrichment_ORF3a int proteins

Supplementary Data 16: Lung_mirnet_enrichment_ORF3a int proteins

## Acknowledgements

Mr. Soura Chakraborty was a recipient of M.Sc., student fellowship from Department of Biotechnology (DBT), Government of India. Ms. Shrabonti Chatterjee is a recipient of Non – National Eligibility Test (Non – NET) fellowship from University Grants Commission (UGC), Government of India. Ms. Subhashree Mardi and Mr. Joydeep Mahata are recipients of DST – INSPIRE (Department of Science and Technology – Innovation in Science Pursuit for Inspired Research) Post – Graduate fellowship from Department of Science and Technology (DST), Government of India. We thank the above funding agencies for their financial support.

## References

1. Dróżdż, M.; Krzyżek, P.; Dudek, B.; Makuch, S.; Janczura, A.; Paluch, E. Current State of Knowledge about Role of Pets in Zoonotic Transmission of SARS-CoV-2. Viruses 2021, 13 (6), 1149. 10.3390/v13061149.

2. Larsen, H. D.; Fonager, J.; Lomholt, F. K.; Dalby, T.; Benedetti, G.; Kristensen, B.; Urth, T. R.; Rasmussen, M.; Lassaunière, R.; Rasmussen, T. B.; Strandbygaard, B.; Lohse, L.; Chaine, M.; Møller, K. L.; Berthelsen, A.-S. N.; Nørgaard, S. K.; Sönksen, U. W.; Boklund, A. E.; Hammer, A. S.; Belsham, G. J.; Krause, T. G.; Mortensen, S.; Bøtner, A.; Fomsgaard, A.; Mølbak, K. Preliminary Report of an Outbreak of SARS-CoV-2 in Mink and Mink Farmers Associated with Community Spread, Denmark, June to November 2020. Euro Surveill 2021, 26 (5). 10.2807/1560-7917.ES.2021.26.5.210009.

3. Meekins, D. A.; Gaudreault, N. N.; Richt, J. A. Natural and Experimental SARS-CoV-2 Infection in Domestic and Wild Animals. Viruses 2021, 13 (10), 1993. 10.3390/v13101993.

4. Prince, T.; Smith, S. L.; Radford, A. D.; Solomon, T.; Hughes, G. L.; Patterson, E. I. SARS-CoV-2 Infections in Animals: Reservoirs for Reverse Zoonosis and Models for Study. Viruses 2021, 13 (3), 494. 10.3390/v13030494.

5. Sharun, K.; Tiwari, R.; Natesan, S.; Dhama, K. SARS-CoV-2 Infection in Farmed Minks, Associated Zoonotic Concerns, and Importance of the One Health Approach during the Ongoing COVID-19 Pandemic. Vet Q 2021, 41 (1), 50–60. 10.1080/01652176.2020.1867776.

6. Eckstrand, C. D.; Baldwin, T. J.; Rood, K. A.; Clayton, M. J.; Lott, J. K.; Wolking, R. M.; Bradway, D. S.; Baszler, T. An Outbreak of SARS-CoV-2 with High Mortality in Mink (Neovison Vison) on Multiple Utah Farms. PLoS Pathog 2021, 17 (11), e1009952. 10.1371/journal.ppat.1009952.

7. Peiris, J. S. M.; Guan, Y.; Yuen, K. Y. Severe Acute Respiratory Syndrome. Nat Med 2004, 10 (Suppl 12), S88–S97. 10.1038/nm1143.

8. Aguanno, R.; ElIdrissi, A.; Elkholy, A. A.; Ben Embarek, P.; Gardner, E.; Grant, R.; Mahrous, H.; Malik, M. R.; Pavade, G.; VonDobschuetz, S.; Wiersma, L.; Van Kerkhove, M. D. MERS: Progress on the Global Response, Remaining Challenges and the Way Forward. Antiviral Research 2018, 159, 35–44. 10.1016/j.antiviral.2018.09.002.

9. Lindner, D.; Fitzek, A.; Bräuninger, H.; Aleshcheva, G.; Edler, C.; Meissner, K.; Scherschel, K.; Kirchhof, P.; Escher, F.; Schultheiss, H.-P.; Blankenberg, S.; Püschel, K.; Westermann, D. Association of Cardiac Infection With SARS-CoV-2 in Confirmed COVID-19 Autopsy Cases. JAMA Cardiol 2020, 5 (11), 1281–1285. 10.1001/jamacardio.2020.3551.

10. Pellegrini, L.; Albecka, A.; Mallery, D. L.; Kellner, M. J.; Paul, D.; Carter, A. P.; James, L. C.; Lancaster, M. A. SARS-CoV-2 Infects the Brain Choroid Plexus and Disrupts the Blood-CSF Barrier in Human Brain Organoids. Cell Stem Cell 2020, 27 (6), 951–961.e5. 10.1016/j.stem.2020.10.001.

11. Scaldaferri, F.; Ianiro, G.; Privitera, G.; Lopetuso, L. R.; Vetrone, L. M.; Petito, V.; Pugliese, D.; Neri, M.; Cammarota, G.; Ringel, Y.; Costamagna, G.; Gasbarrini, A.; Boskoski, I.; Armuzzi, A. The Thrilling Journey of SARS-CoV-2 into the Intestine: From Pathogenesis to Future Clinical Implications. Inflamm Bowel Dis 2020, 26 (9), 1306–1314. 10.1093/ibd/izaa181.

12. Trypsteen, W.; Cleemput, J. V.; Snippenberg, W. van; Gerlo, S.; Vandekerckhove, L. On the Whereabouts of SARS-CoV-2 in the Human Body: A Systematic Review. PLOS Pathogens 2020, 16 (10), e1009037. 10.1371/journal.ppat.1009037.

13. Vijayan, A.; Humphreys, B. D. SARS-CoV-2 in the Kidney: Bystander or Culprit? Nat Rev Nephrol 2020, 16 (12), 703–704. 10.1038/s41581-020-00354-7.

14. Anand, K.; Ziebuhr, J.; Wadhwani, P.; Mesters, J. R.; Hilgenfeld, R. Coronavirus Main Proteinase (3CL ^pro^) Structure: Basis for Design of Anti-SARS Drugs. Science 2003, 300 (5626), 1763–1767. 10.1126/science.1085658.

15. Liu, C.; Zhou, Q.; Li, Y.; Garner, L. V.; Watkins, S. P.; Carter, L. J.; Smoot, J.; Gregg, A. C.; Daniels, A. D.; Jervey, S.; Albaiu, D. Research and Development on Therapeutic Agents and Vaccines for COVID-19 and Related Human Coronavirus Diseases. ACS Cent. Sci. 2020, 6 (3), 315–331. 10.1021/acscentsci.0c00272.

16. Liu, H.; Gai, S.; Wang, X.; Zeng, J.; Sun, C.; Zhao, Y.; Zheng, Z. Single-Cell Analysis of SARS-CoV-2 Receptor ACE2 and Spike Protein Priming Expression of Proteases in the Human Heart. Cardiovasc Res 2020, 116 (10), 1733–1741. 10.1093/cvr/cvaa191.

17. Mifsud, E. J.; Hayden, F. G.; Hurt, A. C. Antivirals Targeting the Polymerase Complex of Influenza Viruses. Antiviral Research 2019, 169, 104545. 10.1016/j.antiviral.2019.104545.

18. Morse, J. S.; Lalonde, T.; Xu, S.; Liu, W. R. Learning from the Past: Possible Urgent Prevention and Treatment Options for Severe Acute Respiratory Infections Caused by 2019- NCoV. ChemBioChem 2020, 21 (5), 730–738. 10.1002/cbic.202000047.

19. Zhou, Y.; Vedantham, P.; Lu, K.; Agudelo, J.; Carrion, R.; Nunneley, J. W.; Barnard, D.; Pöhlmann, S.; McKerrow, J. H.; Renslo, A. R.; Simmons, G. Protease Inhibitors Targeting Coronavirus and Filovirus Entry. Antiviral Research 2015, 116, 76–84. 10.1016/j.antiviral.2015.01.011.

20. Scott, C.; Griffin, S. Viroporins: Structure, Function and Potential as Antiviral Targets. Journal of General Virology 2015, 96 (8), 2000–2027. 10.1099/vir.0.000201.

21. Cady, S. D.; Schmidt-Rohr, K.; Wang, J.; Soto, C. S.; DeGrado, W. F.; Hong, M. Structure of the Amantadine Binding Site of Influenza M2 Proton Channels In Lipid Bilayers. Nature 2010, 463 (7281), 689–692. 10.1038/nature08722.

22. Duff, K. C.; Gilchrist, P. J.; Saxena, A. M.; Bradshaw, J. P. Neutron Diffraction Reveals the Site of Amantadine Blockade in the Influenza A M2 Ion Channel. Virology 1994, 202 (1), 287–293. 10.1006/viro.1994.1345.

23. Hay, A. J.; Wolstenholme, A. J.; Skehel, J. J.; Smith, M. H. The Molecular Basis of the Specific Anti-Influenza Action of Amantadine. The EMBO Journal 1985, 4 (11), 3021–3024. 10.1002/j.1460-2075.1985.tb04038.x.

24. Hong, M.; DeGrado, W. F. Structural Basis for Proton Conduction and Inhibition by the Influenza M2 Protein. Nature 2008, 451 (7178), 596–599. 10.1038/nature06528.

25. Hu, F.; Luo, W.; Cady, S. D.; Hong, M. Conformational Plasticity of the Influenza A M2 Transmembrane Helix in Lipid Bilayers under Varying PH, Drug Binding, and Membrane Thickness. Biochimica et Biophysica Acta (BBA) - Biomembranes 2011, 1808 (1), 415–423. 10.1016/j.bbamem.2010.09.014.

26. Griffin, S. D. C.; Beales, L. P.; Clarke, D. S.; Worsfold, O.; Evans, S. D.; Jaeger, J.; Harris, M.P. G.; Rowlands, D. J. The P7 Protein of Hepatitis C Virus Forms an Ion Channel That Is Blocked by the Antiviral Drug, Amantadine. FEBS Letters 2003, 535 (1–3), 34–38. 10.1016/S0014-5793(02)03851-6.

27. Pavlovic, D.; Neville, D. C. A.; Argaud, O.; Blumberg, B.; Dwek, R. A.; Fischer, W. B.; Zitzmann, N. The Hepatitis C Virus P7 Protein Forms an Ion Channel That Is Inhibited by Long-Alkyl-Chain Iminosugar Derivatives. Proceedings of the National Academy of Sciences 2003, 100 (10), 6104–6108. 10.1073/pnas.1031527100.

28. Premkumar, A.; Wilson, L.; Ewart, G. d; Gage, P. w. Cation-Selective Ion Channels Formed by P7 of Hepatitis C Virus Are Blocked by Hexamethylene Amiloride. FEBS Letters 2004, 557 (1–3), 99–103. 10.1016/S0014-5793(03)01453-4.

29. Foster, T. L.; Verow, M.; Wozniak, A. L.; Bentham, M. J.; Thompson, J.; Atkins, E.; Weinman, S. A.; Fishwick, C.; Foster, R.; Harris, M.; Griffin, S. Resistance Mutations Define Specific Antiviral Effects for Inhibitors of the Hepatitis C Virus P7 Ion Channel. Hepatology 2011, 54 (1), 79–90. 10.1002/hep.24371.

30. Griffin, S.; Stgelais, C.; Owsianka, A. M.; Patel, A. H.; Rowlands, D.; Harris, M. Genotype-Dependent Sensitivity of Hepatitis C Virus to Inhibitors of the P7 Ion Channel. Hepatology 2008, 48 (6), 1779–1790. 10.1002/hep.22555.

31. Shiryaev, V. A.; Radchenko, E. V.; Palyulin, V. A.; Zefirov, N. S.; Bormotov, N. I.; Serova, O. A.; Shishkina, L. N.; Baimuratov, M. R.; Bormasheva, K. M.; Gruzd, Y. A.; Ivleva, E. A.; Leonova, M. V.; Lukashenko, A. V.; Osipov, D. V.; Osyanin, V. A.; Reznikov, A. N.; Shadrikova, V. A.; Sibiryakova, A. E.; Tkachenko, I. M.; Klimochkin, Y. N. Molecular Design, Synthesis and Biological Evaluation of Cage Compound-Based Inhibitors of Hepatitis C Virus P7 Ion Channels. European Journal of Medicinal Chemistry 2018, 158, 214–235. 10.1016/j.ejmech.2018.08.009.

32. Koff, W. C.; Elm, J. L.; Halstead, S. B. Inhibition of Dengue Virus Replication by Amantadine Hydrochloride. Antimicrob Agents Chemother 1980, 18 (1), 125–129. 10.1128/AAC.18.1.125.

33. Lin, C.-C.; Chen, W.-C. Treatment Effectiveness of Amantadine Against Dengue Virus Infection. Am J Case Rep 2016, 17, 921–924. 10.12659/AJCR.901014.

34. Castaño-Rodriguez, C.; Honrubia, J. M.; Gutiérrez-Álvarez, J.; DeDiego, M. L.; Nieto-Torres, J. L.; Jimenez-Guardeño, J. M.; Regla-Nava, J. A.; Fernandez-Delgado, R.; Verdia-Báguena, C.; Queralt-Martín, M.; Kochan, G.; Perlman, S.; Aguilella, V. M.; Sola, I.; Enjuanes, L. Role of Severe Acute Respiratory Syndrome Coronavirus Viroporins E, 3a, and 8a in Replication and Pathogenesis. mBio 2018, 9 (3), e02325-17. 10.1128/mBio.02325-17.

35. Kern, D. M.; Sorum, B.; Mali, S. S.; Hoel, C. M.; Sridharan, S.; Remis, J. P.; Toso, D. B.; Kotecha, A.; Bautista, D. M.; Brohawn, S. G. Cryo-EM Structure of SARS-CoV-2 ORF3a in Lipid Nanodiscs. Nat Struct Mol Biol 2021, 28 (7), 573–582. 10.1038/s41594-021-00619-0.

36. Chen, I.-Y.; Moriyama, M.; Chang, M.-F.; Ichinohe, T. Severe Acute Respiratory Syndrome Coronavirus Viroporin 3a Activates the NLRP3 Inflammasome. Frontiers in Microbiology 2019, 10, 50. 10.3389/fmicb.2019.00050.

37. Farag, N. S.; Breitinger, U.; Breitinger, H. G.; El Azizi, M. A. Viroporins and Inflammasomes: A Key to Understand Virus-Induced Inflammation. The International Journal of Biochemistry & Cell Biology 2020, 122, 105738. 10.1016/j.biocel.2020.105738.

38. Nieto-Torres, J. L.; Verdiá-Báguena, C.; Jimenez-Guardeño, J. M.; Regla-Nava, J. A.; Castaño- Rodriguez, C.; Fernandez-Delgado, R.; Torres, J.; Aguilella, V. M.; Enjuanes, L. Severe Acute Respiratory Syndrome Coronavirus E Protein Transports Calcium Ions and Activates the NLRP3 Inflammasome. Virology 2015, 485, 330–339. 10.1016/j.virol.2015.08.010.

39. Siu, K.-L.; Yuen, K.-S.; Castano-Rodriguez, C.; Ye, Z.-W.; Yeung, M.-L.; Fung, S.-Y.; Yuan, S.; Chan, C.-P.; Yuen, K.-Y.; Enjuanes, L.; Jin, D.-Y. Severe Acute Respiratory Syndrome Coronavirus ORF3a Protein Activates the NLRP3 Inflammasome by Promoting TRAF3- Dependent Ubiquitination of ASC. The FASEB Journal 2019, 33 (8), 8865–8877. 10.1096/fj.201802418R.

40. Qu, Y.; Wang, X.; Zhu, Y.; Wang, W.; Wang, Y.; Hu, G.; Liu, C.; Li, J.; Ren, S.; Xiao, M. Z. X.; Liu, Z.; Wang, C.; Fu, J.; Zhang, Y.; Li, P.; Zhang, R.; Liang, Q. ORF3a-Mediated Incomplete Autophagy Facilitates Severe Acute Respiratory Syndrome Coronavirus-2 Replication. Frontiers in Cell and Developmental Biology 2021, 9, 2012. 10.3389/fcell.2021.716208.

41. Zeng, F.; Huang, Y.; Guo, Y.; Yin, M.; Chen, X.; Xiao, L.; Deng, G. Association of Inflammatory Markers with the Severity of COVID-19: A Meta-Analysis. International Journal of Infectious Diseases 2020, 96, 467–474. 10.1016/j.ijid.2020.05.055.

42. Tan, Y.-J.; Goh, P.-Y.; Fielding, B. C.; Shen, S.; Chou, C.-F.; Fu, J.-L.; Leong, H. N.; Leo, Y. S.; Ooi, E. E.; Ling, A. E.; Lim, S. G.; Hong, W. Profiles of Antibody Responses against Severe Acute Respiratory Syndrome Coronavirus Recombinant Proteins and Their Potential Use as Diagnostic Markers. Clinical and Vaccine Immunology 2004, 11 (2), 362–371. 10.1128/CDLI.11.2.362-371.2004.

43. Zhong, X.; Guo, Z.; Yang, H.; Peng, L.; Xie, Y.; Wong, T.-Y.; Lai, S.-T.; Guo, Z. Amino Terminus of the SARS Coronavirus Protein 3a Elicits Strong, Potentially Protective Humoral Responses in Infected Patients. Journal of General Virology 2006, 87 (2), 369–373. 10.1099/vir.0.81078-0.

44. Liang, T.; Cheng, M.; Teng, F.; Wang, H.; Deng, Y.; Zhang, J.; Qin, C.; Guo, S.; Zhao, H.; Yu, X. Proteome-Wide Epitope Mapping Identifies a Resource of Antibodies for SARS-CoV-2 Detection and Neutralization. Sig Transduct Target Ther 2021, 6 (1), 1–3. 10.1038/s41392-021-00573-9.

45. Wang, H.; Wu, X.; Zhang, X.; Hou, X.; Liang, T.; Wang, D.; Teng, F.; Dai, J.; Duan, H.; Guo, S.; Li, Y.; Yu, X. SARS-CoV-2 Proteome Microarray for Mapping COVID-19 Antibody Interactions at Amino Acid Resolution. ACS Cent. Sci. 2020, 6 (12), 2238–2249. 10.1021/acscentsci.0c00742.

46. Ren, Y.; Shu, T.; Wu, D.; Mu, J.; Wang, C.; Huang, M.; Han, Y.; Zhang, X.-Y.; Zhou, W.; Qiu, Y.; Zhou, X. The ORF3a Protein of SARS-CoV-2 Induces Apoptosis in Cells. Cell Mol Immunol 2020, 17 (8), 881–883. 10.1038/s41423-020-0485-9.

47. Chen, D.; Zheng, Q.; Sun, L.; Ji, M.; Li, Y.; Deng, H.; Zhang, H. ORF3a of SARS-CoV-2 Promotes Lysosomal Exocytosis-Mediated Viral Egress. Dev Cell 2021, 56 (23), 3250–3263.e5. 10.1016/j.devcel.2021.10.006.

48. Miao, G.; Zhao, H.; Li, Y.; Ji, M.; Chen, Y.; Shi, Y.; Bi, Y.; Wang, P.; Zhang, H. ORF3a of the COVID-19 Virus SARS-CoV-2 Blocks HOPS Complex-Mediated Assembly of the SNARE Complex Required for Autolysosome Formation. Dev Cell 2021, 56 (4), 427–442.e5. 10.1016/j.devcel.2020.12.010.

49. Zhang, Y.; Sun, H.; Pei, R.; Mao, B.; Zhao, Z.; Li, H.; Lin, Y.; Lu, K. The SARS-CoV-2 Protein ORF3a Inhibits Fusion of Autophagosomes with Lysosomes. Cell Discov 2021, 7 (1), 1–12. 10.1038/s41421-021-00268-z.

50. Majumdar, P.; Niyogi, S. ORF3a Mutation Associated with Higher Mortality Rate in SARS-CoV-2 Infection. Epidemiol Infect 2020, 148, e262. 10.1017/S0950268820002599.

51. Gordon, D. E.; Jang, G. M.; Bouhaddou, M.; Xu, J.; Obernier, K.; White, K. M.; O’Meara, M.J.; Rezelj, V. V.; Guo, J. Z.; Swaney, D. L.; Tummino, T. A.; Hüttenhain, R.; Kaake, R. M.; Richards, A. L.; Tutuncuoglu, B.; Foussard, H.; Batra, J.; Haas, K.; Modak, M.; Kim, M.; Haas, P.; Polacco, B. J.; Braberg, H.; Fabius, J. M.; Eckhardt, M.; Soucheray, M.; Bennett, M. J.; Cakir, M.; McGregor, M. J.; Li, Q.; Meyer, B.; Roesch, F.; Vallet, T.; Mac Kain, A.; Miorin, L.; Moreno, E.; Naing, Z. Z. C.; Zhou, Y.; Peng, S.; Shi, Y.; Zhang, Z.; Shen, W.; Kirby, I. T.; Melnyk, J. E.; Chorba, J. S.; Lou, K.; Dai, S. A.; Barrio-Hernandez, I.; Memon, D.; Hernandez-Armenta, C.; Lyu, J.; Mathy, C. J. P.; Perica, T.; Pilla, K. B.; Ganesan, S. J.; Saltzberg, D. J.; Rakesh, R.; Liu, X.; Rosenthal, S. B.; Calviello, L.; Venkataramanan, S.; Liboy-Lugo, J.; Lin, Y.; Huang, X.-P.; Liu, Y.; Wankowicz, S. A.; Bohn, M.; Safari, M.; Ugur, F. S.; Koh, C.; Savar, N. S.; Tran, Q. D.; Shengjuler, D.; Fletcher, S. J.; O’Neal, M. C.; Cai, Y.; Chang, J. C. J.; Broadhurst, D. J.; Klippsten, S.; Sharp, P. P.; Wenzell, N. A.; Kuzuoglu-Ozturk, D.; Wang, H.-Y.; Trenker, R.; Young, J. M.; Cavero, D. A.; Hiatt, J.; Roth, T. L.; Rathore, U.; Subramanian, A.; Noack, J.; Hubert, M.; Stroud, R. M.; Frankel, A. D.; Rosenberg, O. S.; Verba, K. A.; Agard, D. A.; Ott, M.; Emerman, M.; Jura, N.; von Zastrow, M.; Verdin, E.; Ashworth, A.; Schwartz, O.; d’Enfert, C.; Mukherjee, S.; Jacobson, M.; Malik, H. S.; Fujimori, D. G.; Ideker, T.; Craik, C. S.; Floor, S. N.; Fraser, J. S.; Gross, J. D.; Sali, A.; Roth, B. L.; Ruggero, D.; Taunton, J.; Kortemme, T.; Beltrao, P.; Vignuzzi, M.; García-Sastre, A.; Shokat, K. M.; Shoichet, B. K.; Krogan, N. J. A SARS-CoV-2 Protein Interaction Map Reveals Targets for Drug Repurposing. Nature 2020, 583 (7816), 459–468. 10.1038/s41586-020-2286-9.

52. Szklarczyk, D.; Kirsch, R.; Koutrouli, M.; Nastou, K.; Mehryary, F.; Hachilif, R.; Gable, A. L.; Fang, T.; Doncheva, N. T.; Pyysalo, S.; Bork, P.; Jensen, L. J.; von Mering, C. The STRING Database in 2023: Protein–Protein Association Networks and Functional Enrichment Analyses for Any Sequenced Genome of Interest. Nucleic Acids Research 2023, 51 (D1), D638–D646. 10.1093/nar/gkac1000.

53. Shannon, P.; Markiel, A.; Ozier, O.; Baliga, N. S.; Wang, J. T.; Ramage, D.; Amin, N.; Schwikowski, B.; Ideker, T. Cytoscape: A Software Environment for Integrated Models of Biomolecular Interaction Networks. Genome Res. 2003, 13 (11), 2498–2504. 10.1101/gr.1239303.

54. Chin, C.-H.; Chen, S.-H.; Wu, H.-H.; Ho, C.-W.; Ko, M.-T.; Lin, C.-Y. CytoHubba:Identifying Hub Objects and Sub-Networks from Complex Interactome. BMC Syst Biol 2014,8 (S4), S11. 10.1186/1752-0509-8-S4-S11.

55. Bader, G. D.; Hogue, C. W. An Automated Method for Finding Molecular Complexes in Large Protein Interaction Networks. BMC Bioinformatics 2003.

56. Maere, S.; Heymans, K.; Kuiper, M. BiNGO: A Cytoscape Plugin to Assess Overrepresentation of Gene Ontology Categories in Biological Networks. Bioinformatics 2005, 21 (16), 3448–3449. 10.1093/bioinformatics/bti551.

57. Kotlyar, M.; Pastrello, C.; Sheahan, N.; Jurisica, I. Integrated Interactions Database: Tissue-Specific View of the Human and Model Organism Interactomes. Nucleic Acids Res 2016, 44 (D1), D536–541. 10.1093/nar/gkv1115.

58. Yu, G.; Wang, L.-G.; Han, Y.; He, Q.-Y. ClusterProfiler: An R Package for Comparing Biological Themes among Gene Clusters. OMICS 2012, 16 (5), 284–287. 10.1089/omi.2011.0118.

59. Chang, L.; Zhou, G.; Soufan, O.; Xia, J. MiRNet 2.0: Network-Based Visual Analytics for MiRNA Functional Analysis and Systems Biology. Nucleic Acids Res 2020, 48 (W1), W244– W251. 10.1093/nar/gkaa467.

60. Farr, R. J.; Rootes, C. L.; Rowntree, L. C.; Nguyen, T. H. O.; Hensen, L.; Kedzierski, L.;Cheng, A. C.; Kedzierska, K.; Au, G. G.; Marsh, G. A.; Vasan, S. S.; Foo, C. H.; Cowled, C.; Stewart, C. R. Altered MicroRNA Expression in COVID-19 Patients Enables Identification of SARS-CoV-2 Infection. PLoS Pathog 2021, 17 (7), e1009759. 10.1371/journal.ppat.1009759.

61. Jacob, F.; Pather, S. R.; Huang, W.-K.; Zhang, F.; Wong, S. Z. H.; Zhou, H.; Cubitt, B.; Fan, W.; Chen, C. Z.; Xu, M.; Pradhan, M.; Zhang, D. Y.; Zheng, W.; Bang, A. G.; Song, H.; Carlos de la Torre, J.; Ming, G. Human Pluripotent Stem Cell-Derived Neural Cells and Brain Organoids Reveal SARS-CoV-2 Neurotropism Predominates in Choroid Plexus Epithelium. Cell Stem Cell 2020, 27 (6), 937–950.e9. 10.1016/j.stem.2020.09.016.

62. Vishnubalaji, R.; Shaath, H.; Alajez, N. M. Protein Coding and Long Noncoding RNA (LncRNA) Transcriptional Landscape in SARS-CoV-2 Infected Bronchial Epithelial Cells Highlight a Role for Interferon and Inflammatory Response. Genes 2020, 11 (7), 760. 10.3390/genes11070760.

63. Delorey, T. M.; Ziegler, C. G. K.; Heimberg, G.; Normand, R.; Yang, Y.; Segerstolpe, Å.; Abbondanza, D.; Fleming, S. J.; Subramanian, A.; Montoro, D. T.; Jagadeesh, K. A.; Dey, K. K.; Sen, P.; Slyper, M.; Pita-Juárez, Y. H.; Phillips, D.; Biermann, J.; Bloom-Ackermann, Z.; Barkas, N.; Ganna, A.; Gomez, J.; Melms, J. C.; Katsyv, I.; Normandin, E.; Naderi, P.; Popov,Y. V.; Raju, S. S.; Niezen, S.; Tsai, L. T.-Y.; Siddle, K. J.; Sud, M.; Tran, V. M.; Vellarikkal, S.K.; Wang, Y.; Amir-Zilberstein, L.; Atri, D. S.; Beechem, J.; Brook, O. R.; Chen, J.; Divakar,P.; Dorceus, P.; Engreitz, J. M.; Essene, A.; Fitzgerald, D. M.; Fropf, R.; Gazal, S.; Gould, J.; Grzyb, J.; Harvey, T.; Hecht, J.; Hether, T.; Jané-Valbuena, J.; Leney-Greene, M.; Ma, H.; McCabe, C.; McLoughlin, D. E.; Miller, E. M.; Muus, C.; Niemi, M.; Padera, R.; Pan, L.;Pant, D.; Pe’er, C.; Pfiffner-Borges, J.; Pinto, C. J.; Plaisted, J.; Reeves, J.; Ross, M.; Rudy,M.; Rueckert, E. H.; Siciliano, M.; Sturm, A.; Todres, E.; Waghray, A.; Warren, S.; Zhang, S.; Zollinger, D. R.; Cosimi, L.; Gupta, R. M.; Hacohen, N.; Hibshoosh, H.; Hide, W.; Price, A. L.; Rajagopal, J.; Tata, P. R.; Riedel, S.; Szabo, G.; Tickle, T. L.; Ellinor, P. T.; Hung, D.; Sabeti, P. C.; Novak, R.; Rogers, R.; Ingber, D. E.; Jiang, Z. G.; Juric, D.; Babadi, M.; Farhi, S.L.; Izar, B.; Stone, J. R.; Vlachos, I. S.; Solomon, I. H.; Ashenberg, O.; Porter, C. B. M.; Li, B.; Shalek, A. K.; Villani, A.-C.; Rozenblatt-Rosen, O.; Regev, A. COVID-19 Tissue Atlases Reveal SARS-CoV-2 Pathology and Cellular Targets. Nature 2021, 595 (7865), 107–113. 10.1038/s41586-021-03570-8.

64. Baindara, P.; Sarker, M. B.; Earhart, A. P.; Mandal, S. M.; Schrum, A. G. NOTCH Signaling in COVID-19: A Central Hub Controlling Genes, Proteins, and Cells That Mediate SARS-CoV-2 Entry, the Inflammatory Response, and Lung Regeneration. Front Cell Infect Microbiol 2022, 12, 928704. 10.3389/fcimb.2022.928704.

65. Breikaa, R. M.; Lilly, B. The Notch Pathway: A Link Between COVID-19 Pathophysiology and Its Cardiovascular Complications. Front Cardiovasc Med 2021, 8, 681948. 10.3389/fcvm.2021.681948.

66. Sciacchitano, S.; Sacconi, A.; De Vitis, C.; Blandino, G.; Piaggio, G.; Salvati, V.; Napoli, C.; Marchetti, P.; Taurelli, B. S.; Coluzzi, F.; Rocco, M.; Vecchione, A.; Anibaldi, P.; Marcolongo, A.; Ciliberto, G.; Mancini, R.; Capalbo, C. H-Ras Gene Takes Part to the Host Immune Response to COVID-19. Cell Death Discov. 2021, 7 (1), 1–3. 10.1038/s41420-021-00541-w.

67. Samimi, N.; Farjam, M.; Klionsky, D. J.; Rezaei, N. The Role of Autophagy in the Pathogenesis of SARS-CoV-2 Infection in Different Cell Types. Autophagy 2022, 18 (7), 1728–1731. 10.1080/15548627.2021.1989150.

68. Tomić, S.; Đokić, J.; Stevanović, D.; Ilić, N.; Gruden-Movsesijan, A.; Dinić, M.; Radojević, D.; Bekić, M.; Mitrović, N.; Tomašević, R.; Mikić, D.; Stojanović, D.; Čolić, M. Reduced Expression of Autophagy Markers and Expansion of Myeloid-Derived Suppressor Cells Correlate With Poor T Cell Response in Severe COVID-19 Patients. Front. Immunol. 2021, 12, 614599. 10.3389/fimmu.2021.614599.

69. Valečka, J.; Almeida, C. R.; Su, B.; Pierre, P.; Gatti, E. Autophagy and MHC-Restricted Antigen Presentation. Molecular Immunology 2018, 99, 163–170. 10.1016/j.molimm.2018.05.009.

70. Su, W.; Yu, X.; Zhou, C. SARS-CoV-2 ORF3a Induces Incomplete Autophagy via the Unfolded Protein Response. Viruses 2021, 13 (12), 2467. 10.3390/v13122467.

71. Koepke, L.; Hirschenberger, M.; Hayn, M.; Kirchhoff, F.; Sparrer, K. M. Manipulation of Autophagy by SARS-CoV-2 Proteins. Autophagy 2021, 17 (9), 2659–2661. 10.1080/15548627.2021.1953847.

72. Gordon, D. E.; Hiatt, J.; Bouhaddou, M.; Rezelj, V. V.; Ulferts, S.; Braberg, H.; Jureka, A. S.; Obernier, K.; Guo, J. Z.; Batra, J.; Kaake, R. M.; Weckstein, A. R.; Owens, T. W.; Gupta, M.;Pourmal, S.; Titus, E. W.; Cakir, M.; Soucheray, M.; McGregor, M.; Cakir, Z.; Jang, G.;O’Meara, M. J.; Tummino, T. A.; Zhang, Z.; Foussard, H.; Rojc, A.; Zhou, Y.; Kuchenov, D.; Hüttenhain, R.; Xu, J.; Eckhardt, M.; Swaney, D. L.; Fabius, J. M.; Ummadi, M.; Tutuncuoglu, B.; Rathore, U.; Modak, M.; Haas, P.; Haas, K. M.; Naing, Z. Z. C.; Pulido, E. H.; Shi, Y.; Barrio-Hernandez, I.; Memon, D.; Petsalaki, E.; Dunham, A.; Marrero, M. C.; Burke, D.; Koh, C.; Vallet, T.; Silvas, J. A.; Azumaya, C. M.; Billesbølle, C.; Brilot, A. F.; Campbell, M. G.; Diallo, A.; Dickinson, M. S.; Diwanji, D.; Herrera, N.; Hoppe, N.; Kratochvil, H. T.; Liu, Y.; Merz, G. E.; Moritz, M.; Nguyen, H. C.; Nowotny, C.; Puchades, C.; Rizo, A. N.; Schulze-Gahmen, U.; Smith, A. M.; Sun, M.; Young, I. D.; Zhao, J.; Asarnow, D.; Biel, J.; Bowen, A.; Braxton, J. R.; Chen, J.; Chio, C. M.; Chio, U. S.; Deshpande, I.; Doan, L.; Faust, B.; Flores, S.; Jin, M.; Kim, K.; Lam, V. L.; Li, F.; Li, J.; Li, Y.-L.; Li, Y.; Liu, X.; Lo, M.; Lopez, K. E.; Melo, A. A.; Moss, F. R.; Nguyen, P.; Paulino, J.; Pawar, K. I.; Peters, J. K.; Pospiech, T. H.; Safari, M.; Sangwan, S.; Schaefer, K.; Thomas, P. V.; Thwin, A. C.; Trenker, R.; Tse, E.; Tsui,T. K. M.; Wang, F.; Whitis, N.; Yu, Z.; Zhang, K.; Zhang, Y.; Zhou, F.; Saltzberg, D.; QCRG Structural Biology Consortium; Hodder, A. J.; Shun-Shion, A. S.; Williams, D. M.; White, K. M.; Rosales, R.; Kehrer, T.; Miorin, L.; Moreno, E.; Patel, A. H.; Rihn, S.; Khalid, M. M.; Vallejo-Gracia, A.; Fozouni, P.; Simoneau, C. R.; Roth, T. L.; Wu, D.; Karim, M. A.; Ghoussaini, M.; Dunham, I.; Berardi, F.; Weigang, S.; Chazal, M.; Park, J.; Logue, J.; McGrath, M.; Weston, S.; Haupt, R.; Hastie, C. J.; Elliott, M.; Brown, F.; Burness, K. A.; Reid, E.; Dorward, M.; Johnson, C.; Wilkinson, S. G.; Geyer, A.; Giesel, D. M.; Baillie, C.; Raggett, S.; Leech, H.; Toth, R.; Goodman, N.; Keough, K. C.; Lind, A. L.; Zoonomia Consortium; Klesh, R. J.; Hemphill, K. R.; Carlson-Stevermer, J.; Oki, J.; Holden, K.; Maures, T.; Pollard, K. S.; Sali, A.; Agard, D. A.; Cheng, Y.; Fraser, J. S.; Frost, A.; Jura, N.; Kortemme, T.; Manglik, A.; Southworth, D. R.; Stroud, R. M.; Alessi, D. R.; Davies, P.; Frieman, M. B.; Ideker, T.; Abate, C.; Jouvenet, N.; Kochs, G.; Shoichet, B.; Ott, M.; Palmarini, M.; Shokat, K. M.; García-Sastre, A.; Rassen, J. A.; Grosse, R.; Rosenberg, O. S.; Verba, K. A.; Basler, C. F.; Vignuzzi, M.; Peden, A. A.; Beltrao, P.; Krogan, N. J. Comparative Host-Coronavirus Protein Interaction Networks Reveal Pan-Viral Disease Mechanisms. Science 2020, 370 (6521), eabe9403. 10.1126/science.abe9403.

73. Zhang, H.; Zheng, H.; Zhu, J.; Dong, Q.; Wang, J.; Fan, H.; Chen, Y.; Zhang, X.; Han, X.; Li, Q.; Lu, J.; Tong, Y.; Chen, Z. Ubiquitin-Modified Proteome of SARS-CoV-2-Infected Host Cells Reveals Insights into Virus–Host Interaction and Pathogenesis. J. Proteome Res. 2021, 20 (5), 2224–2239. 10.1021/acs.jproteome.0c00758.

74. Douen, A.; Panetti, R.; Kaur, P.; Spiers, S.; Akker, E. INVESTIGATION BETWEEN THE ASSOCIATION OF CALCIUM CHANNELS BLOCKERS ON MORTALITY AND INTUBATION IN PATIENTS WITH COVID-19. CHEST 2021, 160 (4), A2503. 10.1016/j.chest.2021.08.013.

75. Mendez, S. R.; Frank, R. C.; Stevenson, E. K.; Chung, M.; Silverman, M. G. Dihydropyridine Calcium Channel Blockers and the Risk of Severe COVID-19. CHEST 2021, 160 (1), 89–93. 10.1016/j.chest.2021.01.073.

76. Semenzato, L.; Botton, J.; Drouin, J.; Baricault, B.; Vabre, C.; Cuenot, F.; Penso, L.; Herlemont, P.; Sbidian, E.; Weill, A.; Dray-Spira, R.; Zureik, M. Antihypertensive Drugs and COVID-19 Risk. Hypertension 2021, 77 (3), 833–842. 10.1161/HYPERTENSIONAHA.120.16314.

77. Camara, A. K. S.; Zhou, Y.; Wen, P.-C.; Tajkhorshid, E.; Kwok, W.-M. Mitochondrial VDAC1: A Key Gatekeeper as Potential Therapeutic Target. Front Physiol 2017, 8, 460. 10.3389/fphys.2017.00460.

78. Han, Q.; Wang, J.; Luo, H.; Li, L.; Lu, X.; Liu, A.; Deng, Y.; Jiang, Y. TMBIM6, a Potential Virus Target Protein Identified by Integrated Multiomics Data Analysis in SARS-CoV-2- Infected Host Cells. Aging 2021, 13 (7), 9160–9185. 10.18632/aging.202718.

79. Vaidyanathan, R.; Reilly, L.; Eckhardt, L. L. Caveolin-3 Microdomain: Arrhythmia Implications for Potassium Inward Rectifier and Cardiac Sodium Channel. Frontiers in Physiology 2018, 9.

80. Straus, M. R.; Bidon, M. K.; Tang, T.; Jaimes, J. A.; Whittaker, G. R.; Daniel, S. Inhibitors of L-Type Calcium Channels Show Therapeutic Potential for Treating SARS-CoV-2 Infections by Preventing Virus Entry and Spread. ACS Infect. Dis. 2021, 7 (10), 2807–2815. 10.1021/acsinfecdis.1c00023.

